# Disrupting ß-catenin dependent Wnt signaling activates an invasive gene program predictive of colon cancer progression

**DOI:** 10.1101/667030

**Authors:** George T. Chen, Delia F. Tifrea, Rabi Murad, Yung Lyou, Ali Mortazavi, Robert A. Edwards, Marian L. Waterman

## Abstract

The recent classification of colon cancer into molecular subtypes revealed that patients with the poorest prognosis harbor tumors with the lowest levels of Wnt signaling. This is contrary to the long-standing understanding that overactive Wnt signaling promotes tumor progression from early initiation stages through to the later stages including invasion and metastasis. Here, we lower the levels of Wnt signaling in colon cancer via interference with two different steps in the pathway that lie upstream or downstream of the effector protein ß-catenin. We find that these Wnt-reduced cancer cells exhibit a more aggressive disease phenotype, including increased mobility in vitro and localized invasion in an orthotopic mouse model. RNA sequencing reveals that interference with Wnt signaling leads to an upregulation of gene programs that favor cell migration and invasion. We identify a set of upregulated genes common among the Wnt perturbations and find that elevated expression of these genes is strongly predictive of poor patient outcomes in early-invasive colon cancer. These genes may have clinical applications as patient biomarkers or new drug targets to be used in concert with existing therapies.

**One Sentence Summary:** Low Wnt Signaling Leads to Invasive Tumor Phenotypes in Colorectal Cancer.

## Introduction

Globally, colorectal cancer (CRC) is one of the leading causes of cancer-related deaths. Although increased screening, changes in lifestyle habits associated with CRC risk, and improvements in treatment have decreased the CRC mortality rate, metastatic disease continues to be a significant challenge to treat. Approximately 50-60% of CRC patients will develop metastatic disease, and the five-year relative survival for these patients is 13.9% (Davis, 2018).

Mutations in the Wnt signaling pathway are a hallmark of CRC, with 80-90% of patients harboring pathway-activating mutations. In normal colon epithelia, the ß-catenin dependent Wnt signaling pathway is important in regulating the renewal of the epithelial layer, which turns over in its entirety every five to seven days. The epithelium consists of a regular repetition of invaginations known as crypts, at the base of which are highly proliferative intestinal stem cells. These cells respond to a high level of Wnt ligand secretion by their neighboring cells to activate the Wnt pathway and drive cell proliferation and self-renewal gene programs through the nuclear localization of ß-catenin and subsequent binding to LEF/TCF transcription factors. The absence of Wnt ligands allows ß-catenin to be targeted for phosphorylation and degradation by the destruction complex, a group of proteins including the scaffold protein Axin2, the tumor suppressor APC, and the kinases CK1 and GSK3 (Gammons & Bienz, 2018). In CRC, the most common mutations target adenomatous polyposis coli (APC), which then limits the cell’s ability to sequester and degrade ß-catenin, allowing ß-catenin to translocate to the nucleus in the absence of a Wnt ligand binding to a Frizzled:LRP receptor complex. Though the general consensus has been that these activating mutations relieve CRC cells from responding to Wnt ligands, recent evidence has shown that Wnt ligands are still capable of modulating Wnt activity, even in APC-mutant cells (Voloshanenko *et al*, 2013; Seshagiri *et al*, 2012).

In addition to tumor growth, it is thought that the overactive Wnt pathway drives disease progression through the latter stages of invasion and metastasis. Some of the findings that support this include a direct link between Wnt regulation of Snail and Twist expression (ten Berge *et al*, 2008), two genes important in epithelial to mesenchymal transition (EMT) – an activity tightly linked to cell migration and invasion. Both genes have been found to be direct transcriptional targets of ß-catenin:LEF/TCF regulatory complexes. Other observations include the fact that the invasive edge of CRC tumors are frequently enriched in nuclear-localized ß- catenin, suggesting that high Wnt pathway activity exists in this region (Brabletz *et al*, 2001; Jung *et al*, 2001). More recently however, several studies have challenged the dogma surrounding Wnt-driven disease progression. For example, one study reported that advanced colon cancers and their metastases have overall decreased Wnt target gene expression relative to early stage tumors (de Sousa E Melo *et al*, 2011). Though nuclear ß-catenin is observed in these patients’ tumors, de Sousa E Melo *et al* found that several Wnt target genes, including *ASCL2* and *LGR5*, are methylated in advanced disease samples. Additionally, a different group showed that repression of Wnt signaling through expression of dominant negative TCF4 in colon cancer cells increased their metastatic potential when introduced into mice via tail vein injection (Varnat *et al*, 2010). But perhaps the most compelling evidence thus far is that transcriptional signatures of advanced, poor prognosis colon cancers feature decreases in Wnt signaling - a finding that came from a recent effort to classify the tumors from colon cancer patients by molecular subtype (Guinney *et al*, 2015). Combining data from six previous studies that collectively analyzed over 3000 patient tumors and associated clinical data, a four-cluster categorization scheme was developed. Of these four groups, the tumors with the highest Wnt gene signature (labeled CMS2, or Canonical) exhibited the best overall clinical outcome, while the patients with most aggressive disease, the poorest prognosis (CMS4, or mesenchymal) had the lowest Wnt signature expression. CMS4 additionally has a marked upregulation of gene programs associated with matrix remodeling, EMT activation, and stromal infiltration. As a follow-up on this finding, Smedt *et al* analyzed patient samples of both total tumor and portions of small budding cell clusters at the invasive edge and found that the invasive clusters demonstrated a shift from CMS2 to CMS4 – meaning, a shift to an overall lower level of Wnt target gene expression (Smedt *et al*, 2016). The invasive cells additionally harbored upregulation of gene programs connected to EMT, cell movement, cell morphology, and cell survival - and downregulation of programs for cell growth and proliferation. In a following study that aligned PDX models and a large panel of colon cancer cell lines to the CMS system, a clear difference in response to chemotherapy emerged between CMS2 and CMS4 samples; chemotherapy induced apoptosis in CMS2 samples, but CMS4 samples were largely resistant (Linnekamp *et al*, 2018). Interestingly, the researchers were also able to classify a majority of the traditional colon cancer cell lines tested into different CMS groups, even though these cells have been maintained in an *in vitro* system for decades. This suggests that there is at least some level of intrinsic stability of gene expression in cancer cell lines that preserves CMS status *ex vivo*. And finally, a second study to classify colorectal cancers using PDX tumors devoid of human stromal cells identified five CRC/CRIS subtypes (Isella *et al*, 2017). Consistent with the CMS classification, subtypes with the strongest Wnt signaling (CRIS-C,-D,-E) had better prognosis than the subtypes with lower Wnt signaling activity (CRIS-A, -B).

Collectively, the above studies suggest that colorectal cancers with moderate-to-low Wnt signaling have one of the most aggressive tumor phenotypes and the poorest prognosis. Here we directly test this implied correlation by genetically manipulating Wnt/ß-catenin activity in colorectal cancer cells and evaluating their tumor phenotypes at an orthotopic site in the mouse colon. We primarily utilize two colon cancer cell lines, SW480 and SW620. These lines were derived from the same patient, the former from the primary tumor and the latter from a metastatic node. In line with the CMS2 to CMS4 observation described above, the baseline Wnt signaling level of SW620 cells is much lower than that of the SW480 line. We downregulate Wnt signaling activity in multiple ways, and utilizing an orthotopic mouse model, we observe an increase in tumor invasion in all cases. Comparing the differentially expressed genes between the parental and downregulated lines, we directly connect decreases in Wnt signaling with increases in gene expression profiles that promote tumor cell invasion. We demonstrate in multiple patient datasets that overexpression of specific genes is significantly correlated with poor prognosis. We conclude that inhibition of Wnt signaling in Wnt-high colon tumors will lead to a dramatic increase in tumor invasion and aggression.

## Results

### Decreasing Wnt signaling in SW480 cells increases cell invasiveness

Our previous studies determined how decreases in Wnt signaling affect tumor progression in a subcutaneous xenograft tumor model (Pate *et al*, 2014). To decrease Wnt signaling, we lentivirally transduced an expression vector for dominant negative LEF1 (dnLEF1) into SW480 colon cancer cells prior to subcutaneous injection (Figure 1A-C). LEF1 is a LEF/TCF transcription factor that occupies Wnt Response Elements (WREs) in enhancers and promoters and recruits ß-catenin to activate transcription of Wnt target genes. The dnLEF1 expression construct lacks coding sequences for the N-terminal ß-catenin binding domain, but retains all other sequences, including the HMG-box DNA binding domain and nuclear localization signal. This truncated transcription factor can localize to the nucleus and displace endogenous full-length LEF/TCFs from their occupancy of Wnt Response Elements (WREs) at bona fide Wnt target genes. We used this mode of interference as it had been first developed by Van de Wetering and Clevers to effectively interfere with canonical Wnt signaling in the nucleus (Van de Wetering *et al*, 2002). Indeed, using a Wnt-luciferase assay, we find that dnLEF1 transduction-expression decreases Wnt signaling activity by 90% compared to Mock-transduced cells in SW480 (Figure 1I). Three weeks after subcutaneous injection of transduced cells, we harvested the tumors and examined them histologically. We observed that tumors expressing dnLEF1 exhibited a significant decrease of in tumor weight and volume (four to five fold) and a 50% decrease in angiogenic vasculature (Figure 1A-C)(Pate *et al*, 2014). These data are in line with the overall understanding that high levels of Wnt signaling promote tumor growth and development. However, the subcutaneous microenvironment on the back flank of a mouse is not the endogenous location for colon tumor development and further, the subcutaneous environment lacks the nutrient availability of the richly vascularized intestinal epithelium. We therefore expanded our animal studies to include an orthotopic injection model whereby tumors were developed in the mouse colon.

**Figure 1.**
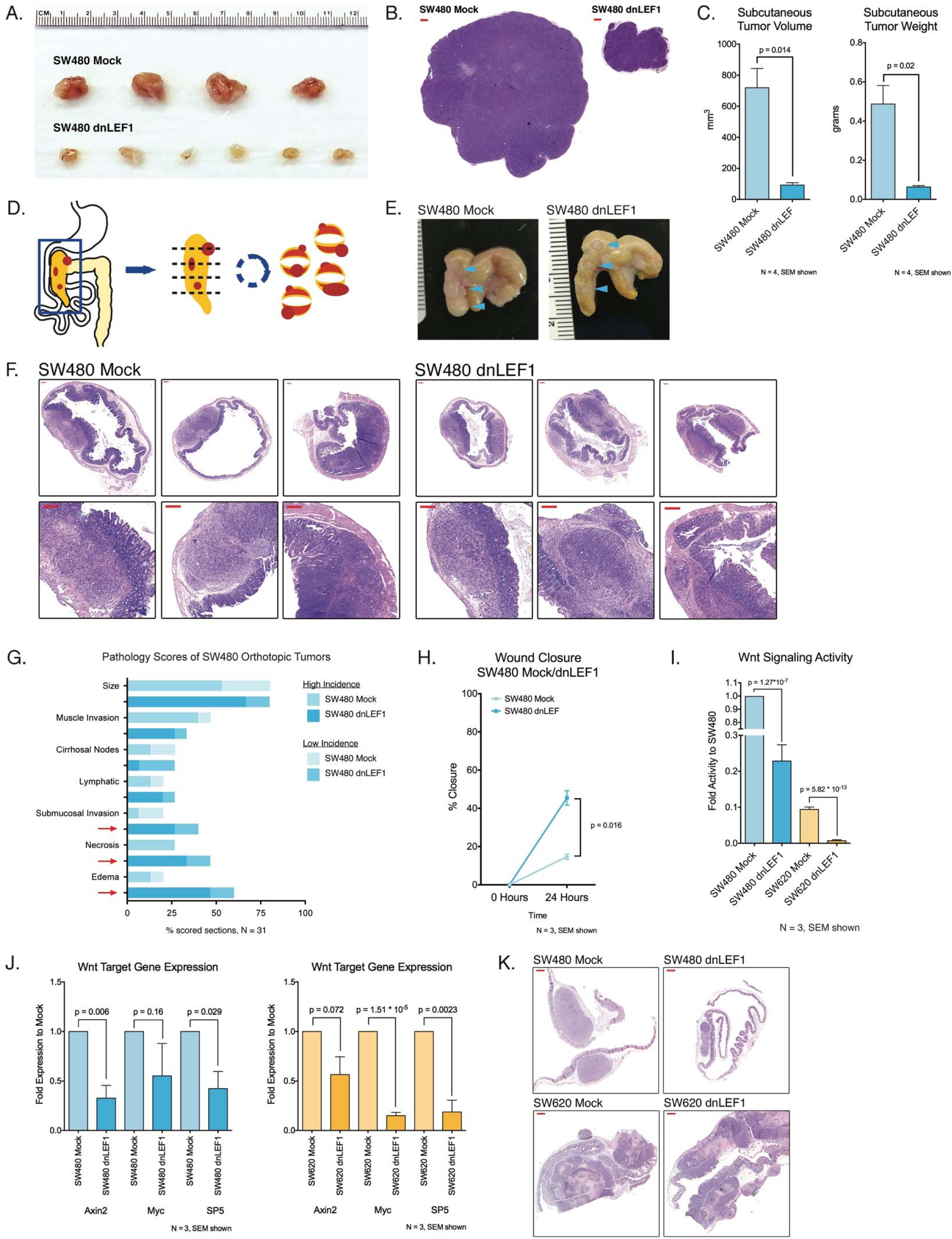
Decreasing Wnt in SW480 colon cancer xenografts increases cell migration and invasion. A-C. dnLEF1-transduced SW480 cells show a significant decrease in tumor burden compared to Mock-transduced SW480 cells in subcutaneous tumors in immunocompromised mice. Representative tumors shown at harvest (A), FFPE sections stained with hematoxylin & eosin (B). Scale bar is 200 um. Quantitation of both tumor volume and weight show statistically significant differences between SW480 Mock and SW480 dnLEF1 (C). D. Schematic of orthotopic injection and harvest. Tumors are injected into the caecum wall and allowed to grow for 3-5 weeks. At harvest, the caecum is cut latidudinally and mounted on edge. E-F. Representative images of caecums at harvest (E) and FFPE sections stained with hematoxylin and eosin (F). Top row: full caecum section, bottom row: magnified inset. All scale bars 200um. G. Blinded pathologist scoring of orthotopic tumor sections shows a greater number of sections ranked high in submucosal invasion, necrosis, and edema in SW480 dnLEF1 tumors compared to SW480 Mock. H. Scratch assay of SW480 Mock and SW480 dnLEF cells on plastic showed that dnLEF1 expression significantly increases cell migration after 24 hours of culture. I. SW620 colon cancer cells inherently exhibit significantly decreased Wnt signaling activity compared to SW480 cells as measured by SuperTOPFlash luciferase activity. Wnt signaling activity is significantly decreased by dnLEF1 expression in both SW480 and SW620 cell lines. J. Quantitative PCR of Wnt target genes *AXIN2*, *MYC*, and *SP5* in SW480 and SW620 cell lines shows significant decrease with dnLEF1 expression. K. Hematoxylin and eosin stains of orthotopic tumor sections. SW480 orthotopic tumors are less aggressive than SW620 orthotopic tumors, expression of dnLEF1 enhances invasion. Scalebars are 500um.

To study tumorigenesis in a more representative microenvironment, we injected the SW480 Mock or SW480 dnLEF1 cells into the stroma between the epithelial and muscle wall layers of the caecum (Figure 1D-E). After four weeks of development, the caecums were harvested and sectioned for immunohistochemistry. Unlike our findings with subcutaneous tumors, we observed no gross deficit in tumor size when comparing the SW480 Mock and SW480 dnLEF1 orthotopic tumors, implying that the high-density vasculature in the colon wall was able to compensate for the deficits in angiogenic development in the subcutaneous-dnLEF1 tumors (Figure 1F). A blinded evaluation of physiological features for each tumor by pathologists revealed that SW480 dnLEF1 orthotopic tumors exhibit higher rates of invasion into the submucosa, higher prevalence of intratumor necrosis, and much more edema (Figure 1G, full images with key in Supplemental Figure 1). When each component of the caecum and tumor was quantified by area on a per-section basis, there was no significant difference in amount of caecum space filled by tumor, but we observed a significant increase in edema induced by the SW480 dnLEF1 tumors. Along with a near-significant difference in the degree of intact epithelium, these results point to a more aggressive tumor formed by SW480 dnLEF1 cells, extravasating and clogging the lymphatic network, and likely leading to the increase in edema (Supplemental Figure 2). These data very clearly suggest that SW480 dnLEF1 tumors are more invasive and their cells more migratory. To evaluate the intrinsic migratory potential of the cells, we plated parental and dnLEF1 expressing cells at high density in a tissue culture plates and wounded the cell monolayer by introducing a “scratch-wound” track. After 24 hours, we observed that SW480 dnLEF1 cells were significantly more migratory into the scratch-wound than SW480 Mock cells (Figure 1H). We conclude that the decrease of Wnt signaling in these colon cancer cells creates a more aggressive tumor both because the cells are intrinsically more migratory and because given the correct microenvironmental context they exhibit invasive behaviors.

The SW480 cell line was derived from the primary tumor of a CRC patient, and a metastatic lymph node from the same patient was used to derive the line SW620. It is known that SW620 cells have significantly less Wnt activity than SW480 (e.g. SW620 Mock is closer in Wnt activity levels to SW480 dnLEF1), and we were interested in the effect of further reducing Wnt signaling in the already Wnt-low line. The addition of the dnLEF1 construct in SW620 cells further significantly reduced Wnt signaling levels (Figure 1I). For both cell lines, expression of dnLEF1 significantly decreases expression of Wnt targets *AXIN2*, *MYC*, and *SP5* (Figure 1J). Unlike SW480, in the orthotopic condition, SW620 Mock cells are already inherently invasive, however, SW620 dnLEF1 appears to still be more aggressive (Figure 1K). Blinded evaluation of SW620 orthotopic tumors showed significant increases in the incidence of extracolonic tumors in SW620 dnLEF1 cells compared to SW620 Mock (Figure 2A-B). In both SW620 Mock and SW620 dnLEF1 tumors, more sections contained necrotic areas compared to SW480 Mock or SW480 dnLEF1 suggesting an innate, higher sensitivity of the SW620 cells to nutrient levels in the surrounding microenvironment. Additional sections revealed an increase in tissue edema, a likely result of tumor cells filling and plugging the lymphatic network to prevent drainage. Using CD298 as a marker of human cells, we used Flow cytometry analysis to detect human cancer cells that had migrated to neighboring mesenteric tissue (Lawson *et al*, 2015). This analysis showed that a significantly higher percentage of SW620 dnLEF1 cells were present in the mesentery of orthotopically-injected mice compared to SW620 Mock cells, indicating a higher level of intravasation into the surrounding lymphatic network (Figure 2B). A “scratch-wound” assay for intrinsic migratory potential also showed that SW620 dnLEF1 cells were significantly more migratory than SW620 Mock cells (Figure 2C). These data suggest that even in cell lines with low Wnt signaling activity, the effect of further decreasing Wnt activity creates a more aggressive tumor.

**Figure 2.**
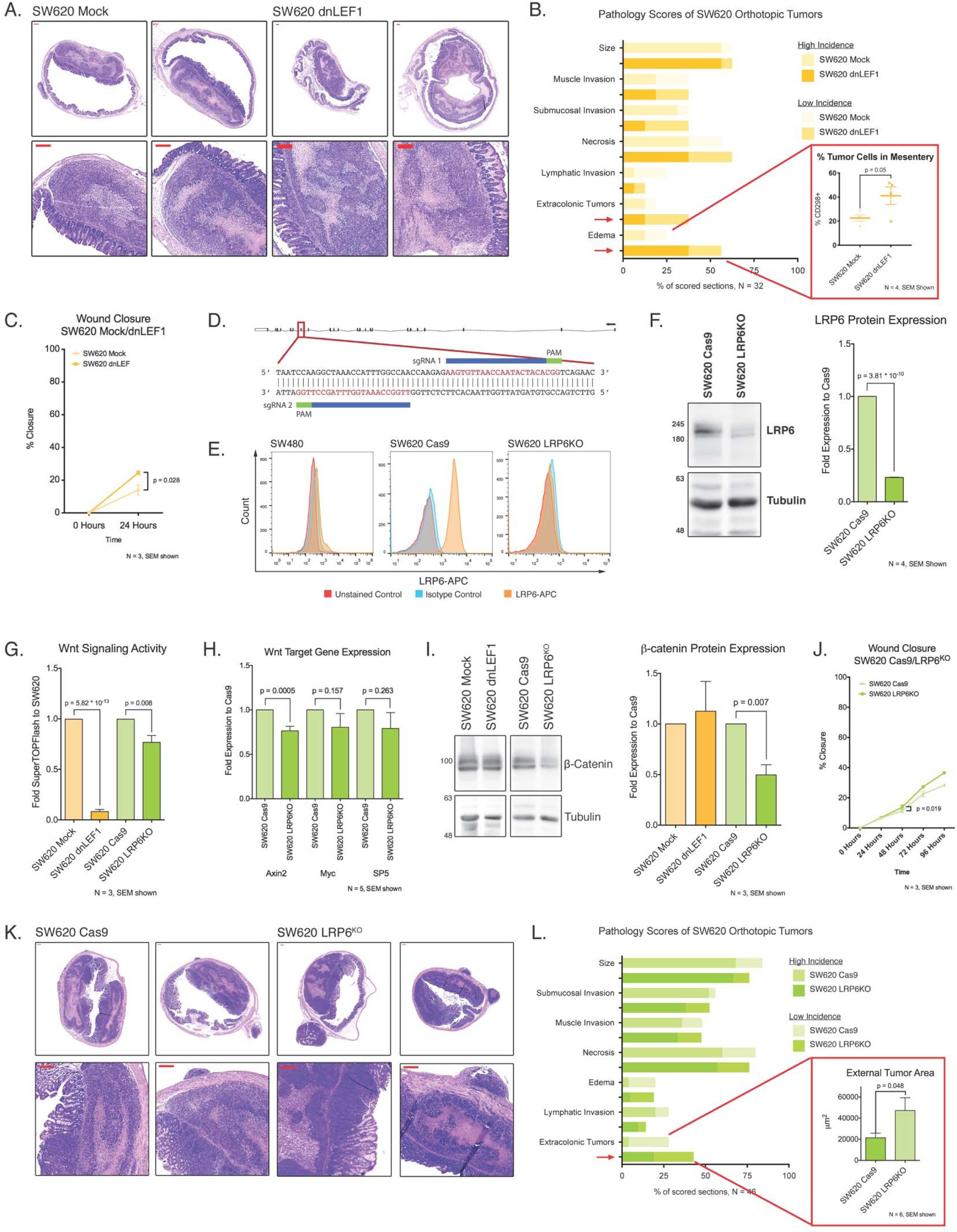
Modulating Wnt signaling upstream or downstream of b-catenin in SW620 cells increases invasion in orthotopic tumors. A. Representative hematoxylin & eosin images of SW620 Mock and SW620 dnLEF1 orthotopic tumors. Scalebars are 200um. B. Blinded pathologist scoring of SW620 Mock and dnLEF1 orthotopic tumor sections. A greater number of dnLEF1tumor sections ranked “high” or “low” for extracolonic tumors and edema compared to Mock tumor sections. Flow cytometry of live cells isolated from the mesentery of orthotopically-injected mice reveals a significant increase of tumor cells detected in the mesentery of SW480 dnLEF1-injected mice. C. Scratch assay of SW620 Mock versus dnLEF1 shows significant difference in cell motility by 24 hours post-scratch. D. Schematic for LRP6 CRISPR effects on b-catenin dependent Wnt signaling. Two CRISPR guide RNAs were developed to target LRP6, their alignments to the nucleotide sequence along with the PAM sites are highlighted in the blue box. Both guide RNAs were used with a Cas9 nickase for specific targeting of LRP6. E. Live cell flow cytometry reveals minimal expression of LRP6 in SW480 cells at the cell membrane, but is visible when cells are fixed (data in supplement). In contrast, SW620 cells robustly express LRP6, and LRP6 CRISPR expression, coupled with FACS sorting and clonal expansion, successfully eliminated LRP6 protein from the cell membrane. F. Validation of LRP6 knockout in SW620 by Western blotting shows a significant decrease in total protein level. G. Wnt signaling levels as measured by SuperTOPFlash luciferase reporter activity, normalized to SW620 Mock. Expression of dnLEF1 reduced Wnt activity by 80-90%. LRP6 knockout in SW620 cells induced a modest, but statistically significant decrease in Wnt signaling activity. H. Expression of Wnt target genes *AXIN2*, *MYC*, and *SP5* in SW620 LRP6KO normalized to parental control. *AXIN2* is significantly decreased; *MYC* and *SP5* trend downwards but not to significance. I. Total ß-catenin protein level is significantly decreased in SW620 LRP6KO, but not SW620 dnLEF1. The loss of LRP6 affects the stability of ß-catenin due to the release of the destruction complex from the plasma membrane. (Saito-diaz *et al*, 2018) J. Scratch assay of SW620 Cas9 versus LRP6KO shows a significant difference in cell motility only after 48 hours, suggesting with a more dramatic Wnt inhibition, there is an increase in cell motility. K. Representative hematoxylin & eosin stained sections of SW620 Cas9 and SW620 LRP6KO orthotopic tumors. Scalebars are 200um. L. Blinded pathologist scoring of SW620 Cas9 and SW620 LRP6KO orthotopic tumor sections. A greater number of LRP6KO sections had extracolonic tumors compared to Cas9 sections. Quantification of extracolonic tumors per section shows no difference in number of extracolonic tumors between SW620 Cas9 and SW620 LRP6KO. Area measurements of extracolonic tumors show a significantly higher average tumor area in LRP6KO samples.

### LRP6 is required for ß-catenin-dependent Wnt signaling in colon cancer cells

Because several groups have reported that Wnt ligands are capable of enhancing Wnt signaling activity, even in APC-mutant colon cancers (Voloshanenko *et al*, 2013; Seshagiri *et al*, 2012; Giannakis *et al*, 2014; Jung *et al*, 2015; Nishioka *et al*, 2011), we hypothesized that autocrine-acting Wnt ligands from the cancer cells and paracrine Wnt signals from the surrounding microenvironment might contribute to maintaining colon cancer cells in a non-invasive state in the orthotopic condition. The dominant negative LEF1 constructs do not directly address this hypothesis, as their effect is to interfere with ß-catenin actions in the nucleus, not the cell membrane. Therefore, to more directly probe how Wnt ligands in the tumor microenvironment affect ß-catenin-dependent signaling and colon cancer cell behavior, we used a CRISPR/Cas9 system to genetically delete the essential Wnt ligand co-receptors LRP5 (Supplemental Figure 3A) and LRP6 (Figure 2D). We tested the efficacy of this manipulation by transiently co-transfecting HEK293 cells with the Cas9 nickase, LRP5 or LRP6 guide RNA expression constructs, and various Wnt ligand expression plasmids for ligands known to be expressed in colon cancer. Using the SuperTOPFlash luciferase reporter, we assayed for Wnt signaling activity. Knockdown of LRP5 or LRP6 expression effectively interfered with Wnt ligand activation of the luciferase reporter. For example, we noted that the Wnt1 ligand required both LRP5 and LRP6 for reporter activation (Supplemental Figure 3), in line with observations from other groups (Goel *et al*, 2012). We tested other Wnt ligands and found that while a number of Wnt ligands required both LRP5 and LRP6, others required solely LRP6, while none of the ligands required only LRP5 (Supplemental Figure 3). In part, this may be due to LRP6 influencing LRP5 expression, as the knockout of LRP6 also reduced LRP5 protein (Supplemental Figure 4F-I). To confirm that this was not an off-target effect of our chosen guide RNAs, we also silenced LRP6 using an shRNA construct, which yielded the same result (Supplemental Figure 4G,I). Given that the LRP6 co-receptor appears to be a uniformly required co-receptor for ß-catenin-dependent signaling, we focused our remaining efforts using LRP6 knockout cells.

### LRP6 modulates Wnt signaling levels in APC-mutant colon cancer

We noted that SW480 cells do not innately express much LRP6 protein and that what residual LRP6 protein is detectable, is not located on the cell membrane where Frizzled receptors reside (Figure 2E, Supplemental Figure 4A-C). We therefore focused our LRP6 knockout efforts on SW620 cells because LRP6 is robustly expressed on the cell surface. Using flow cytometry, we isolated a population of LRP6 knockout cells from the total transfected SW620 cell population (Figure 2E). We validated that LRP6 expression is strongly decreased in SW620 LRP6KO cells by Western blot (Figure 2F). Interestingly, Wnt signaling is decreased by ten to twenty percent compared to expression of Cas9 alone (Figure 2G). However, as expected, compared to the dominant negative LEF1-expressing lines, the effect of a genetic LRP6 knockout on overall Wnt signaling in SW620 cells is not as strong (Figure 2G & 2H). These results demonstrate that decreasing expression of LRP6 moderately decreases Wnt signaling, pointing to real possibilities for ligand-based Wnt pathway activation, even in the presence of a downstream APC mutation. Additionally, we found that ß-catenin protein levels significantly decreased when *LRP6* was deleted, but not with dnLEF1 expression (Figure 2I). While this can be explained in part by a decrease in ß-catenin (*CTNNB1*) mRNA expression (data not shown), the total cellular protein level of ß-catenin decreased more than its *CTNNB1* mRNA, suggesting that ß-catenin protein degradation is enhanced. This is in line with findings from Saito-Diaz *et al* (Saito-Diaz *et al*, 2018), which suggest that LRP6 associates with the ß-catenin destruction complex in the absence of a Wnt ligand, partially inhibiting its activity. Thus the loss of LRP6 allows the destruction complex to be more active, decreasing overall ß-catenin protein. We confirmed this by decreasing LRP6 expression in colon cancer cell line HCT116, which has a mutant ß-catenin and wildtype APC. LRP6 knockout did not significantly decrease Wnt signaling activity in these cells, suggesting that the release of the destruction complex from the plasma membrane did not increase ß-catenin degradation (Supplemental Figure 5A).

Interestingly, when we decreased LRP5 or LRP6 expression in colon cancer cell line COLO320, which has a highly truncated APC, we find that Wnt signaling is reduced by 40-50%. However, ß-catenin expression is not significantly reduced in LRP6-diminished cells, due to the absence of APC domains that interact with LRP6 (Supplemental Figure 5B-D). We conclude that knocking out LRP6 in APC-mutant colon cancer cell lines destabilizes ß-catenin and moderately decreases Wnt signaling.

Orthotopic tumors of SW620 Cas9 and SW620 LRP6KO were largely indistinguishable from one another by pathology scoring, except for again, a higher prevalence of large extracolonic tumors in LRP6KO sections (Figure 2K, L and Supplemental Figures 1 & 2). While the number of extracolonic tumors per section was similar between the two tumor types, the size of the LRP6KO tumors were dramatically larger than the Cas9 tumors (Figure 2L), suggesting that the SW620 LRP6KO have an improved ability to migrate, survive, and/or proliferate outside the colon. To note: subcutaneous SW620 LRP6KO tumors were not significantly smaller than the SW620 Cas9 controls (Supplemental Figure 6). This does not appear to be the case in SW620 dnLEF1 orthotopic tumors, which grow to fill the epithelial space significantly more than SW620 Mock, but do not produce significantly larger extracolonic tumors (Supplemental Figure 2). *In vitro*, SW620 LRP6KO cells were more migratory than SW620 Cas9, but this enhanced phenotype was only evident after 48 hours (Figure 2J). This suggests that there may be a graded effect to Wnt signaling inhibition of cell migration, with decreasing Wnt signaling levels yielding a graded increase in the rate of migration. Using immunohistochemistry, we probed orthotopic tumors for phosphorylated Histone H3 as a marker of proliferation and found no measurable difference between SW620 Cas9 and SW620 LRP6KO orthotopic tumors (Data not shown). Therefore, we conclude that the larger extracolonic tumors from SW620 LRP6KO cells are more likely the result of increased survival and early migration activities rather than increased proliferation.

### RNA Sequencing reveals a common invasive gene signature for decreased ß-catenin-dependent Wnt signaling

To identify changes in gene expression that occur in the conversion to an invasive phenotype, we performed RNAseq of bulk mRNA from the orthotopic tumors, taking advantage of the mixed mouse and human cell environment to analyze concurrent gene expression changes in the human tumor versus the mouse stroma. We also used the relative percentage of reads mapped to human versus mouse genomes to serve as a proxy for assessing the relative abundance of human versus mouse tissue (Figure 3A). We found that the Wnt-low, SW620 tumor cells, both control and further Wnt-diminished, comprised a higher percentage of the tissue samples than the Wnt-high SW480 tumors, with PBS-injected caecums serving as a negative control. Differential gene expression analysis revealed approximately 350 significantly changed genes in SW480 dnLEF1 cells compared to SW480 Mock, but twice as many significant changes in the SW620 Mock versus SW620 dnLEF1. We hypothesize that the greater number of differentially expressed genes in SW620 tumors derives from a more plastic state in SW620 compared to SW480 cells, a state that is more sensitive and responsive to genetic changes. As SW620 cells were derived from a more advanced, metastatic site (lymph node) compared to the primary-tumor derived SW480 line, these cells might have already proceeded through an epithelial-to-mesenchymal transition (EMT), a process that requires de-differentiation. In fact, poorly differentiated tumors are known to be more invasive than well-differentiated tumors, and as such, this state correlates with poor patient prognosis. We compared SW480 Mock tumors to SW620 Mock tumors and found sharp differences between the two cell lines, with over 7000 genes significantly changed in expression. Of these significant gene changes, many are shared with the significantly changed genes between SW480 dnLEF1 and SW480 Mock, suggesting these gene expression changes are associated with Wnt signaling reduction (Sup. Data S1). Not surprisingly, the more moderate change in Wnt signaling induced by knocking out LRP6 led to fewer and smaller, but still-significant changes in gene expression in SW620 (Figure 3B). Because of these minimal changes, we focused our downstream analysis on the dominant negative LEF1-expressing cells.

**Figure 3.**
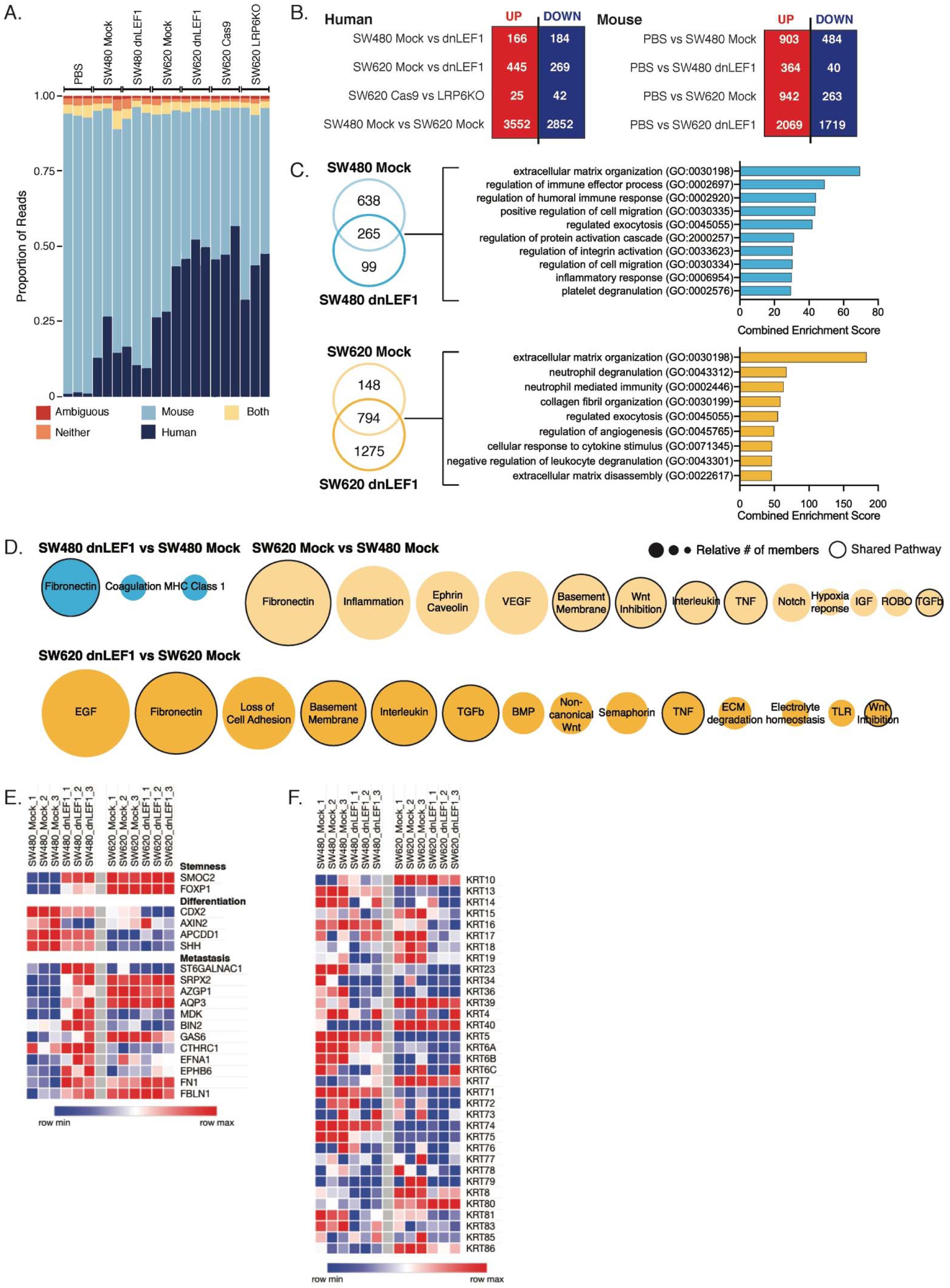
RNA sequencing of orthotopic tumors reveals metastasis-promoting gene expression changes in tumor cells and stromal cells with decreasing Wnt signaling levels. A. Proportion of reads from orthotopic tumor samples mapped to mouse or human genomes. Proportion of human to mouse reads can be considered as proxy for proportion of tissue from each species in each sample. B. Differentially expressed genes reaching significance of p > 0.1 in comparing parental tumors to Wnt-reduced tumors in human genome, and tumor to no tumor in mouse genome. C. Differentially upregulated genes from the mouse stroma conserved between parental and dnLEF1-expressing tumors reveals enrichment in programs affecting extracellular matrix organization, immune response, and cell migration. D. Differentially expressed ligands and receptors expressed by human and mouse genomes reveal shared upregulation in fibronectin signaling in all comparisons. Comparing the more aggressive SW620 parental tumor to SW480 parental tumor reveals upregulation of genes involved in basement membrane signaling, Wnt inhibition, TGFß signaling, TNF signaling, and immune signaling that are similarly changed in comparing SW620 dnLEF1 to SW620 Mock, indicating an increase in signaling with decreasing Wnt. Circles are scaled to indicate the relative number of genes changed per signaling network. Outlined circles highlight signaling networks shared between comparisons. E. Heatmap of selected genes associated with stemness, differentiation, and metastasis shows with decreasing Wnt signaling there is an increase in stemness, decrease in differentiation, and increase in metastasis-related gene expression. F. Heatmap of keratins shows a global trend towards decreased expression with decreased Wnt signaling.

By mapping RNAseq reads to the mouse genome, we compared the stromal signature of each tumor type to the PBS-injected control as a way to identify gene expression changes in the stroma that occur in response to tumor growth (Figure 3B). Both SW480 Mock and SW480 dnLEF1 tumors triggered increases in the expression of 265 genes– genes that are increased in expression no matter what type of tumor is developing in the caecum. Enrichr analysis revealed these genes to be associated with extracellular matrix organization, immune response, and cell migration (Figure 3C) – genes involved in wound repair. The same analysis performed with SW620 tumor data (Mock vs. dnLEF1 and/or LRP6KO) revealed an even greater degree of enrichment in wounding programs of extracellular matrix organization and immune response terms (Figure 3C).

Since communication between cancer cells and their environment affects tumor progression, we sought to identify changes in the expression of receptors and ligands that increase in the invasive tumors. We filtered the list of differentially expressed human genes between SW480 Mock and SW480 dnLEF1 tumors for significantly upregulated ligands and receptors. We added the list of ligands and receptors that were selectively and significantly upregulated in the *mouse* stroma in the SW480 dnLEF1 orthotopic tumors (above and beyond the ligands and receptors upregulated in SW480 Mock tumors). The resulting list was subjected to network analysis using GeNets (Li *et al*, 2018). This process was repeated comparing SW480 Mock versus SW620 Mock and SW620 Mock versus SW620 dnLEF1, illustrating a sequential change from highest level of Wnt signaling to lowest. We identify highly conserved networks that are increasingly upregulated in invasive tumors. Conserved between the three network comparisons is the strong presence of fibronectin signaling, which has been previously shown to play a role in cell invasion (Hanamura *et al*, 1997)(Figure 3D). In the more aggressive tumors, additional changes in signaling to the basement membrane, inhibiting Wnt signaling, and activating TGFβ signaling are conserved. As these networks were derived from uniquely upregulated ligands and receptors in the stroma, the sharing of pathways between tumor-set comparisons suggests that there is significant upregulation at each progressive level of decreased Wnt signaling. This data therefore underscores how Wnt-active cancer cells might influence the cellular communication to stromal cells to moderate and suppress its own invasive phenotype. This analysis also shows that expression of extracellular matrix organization genes is not enough to induce cell migration for, as these genes are already upregulated in the stroma of the non-invasive SW480 Mock tumors.

Of the significantly upregulated genes in the invasive orthotopic tumors, we find that among the top changes are genes involved in stemness and metastasis. Taken together, these changes suggest that with a decrease of Wnt signaling, there is an increase in pluripotency-related gene expression, a decrease in differentiation-related expression, and an increase in metastasis-related expression (Figure 3E). Among the most significantly downregulated genes, we consistently observe that the expression of many keratins decrease in parallel with decreasing Wnt activity (Figure 3F), indicating a loss of epithelial identity.

### Decreased Wnt signaling in colorectal cancer patients is correlated with poor prognosis

Recent attempts to categorize colorectal cancer patients by molecular biomarkers have consistently shown that patients with the highest levels of Wnt signaling have the best prognosis, and patients with the lowest levels have the poorest (Guinney *et al*, 2015; Isella *et al*, 2017). To determine if our observations with decreased Wnt signaling in orthotopic tumors translated to the patient experience., we analyzed gene expression data from rectal cancer patients (GDS3756, (Snipstad *et al*, 2010)). We determined that there are significant decreases in Wnt target gene expression when patients are treated with radiochemotherapy (Figure 4A; *AXIN2*, *LEF1*, and *SP5*) suggesting that the actual activity of the pathway is suppressed. Interestingly, there is consistent upregulation of *GAS6*, *FN1*, *FBLN1*, and *SRPX2* expression in treated patients – changes that closely parallel increased expression in invasive orthotopic tumors (Figure 4B). GAS6 is a secreted ligand that binds to AXL, a well-characterized tyrosine kinase receptor that activates cell migration signaling (Axelrod & Pienta, 2014; Goyette *et al*, 2018). FN1 is known to interact with FBLN1 in the extracellular matrix and has been implicated in promoting cell migration (Pupa *et al*, 2002). SRPX2 is a ligand for the urokinase plasminogen activator surface receptor and has been shown to be overexpressed in gastrointestinal cancers and to promote cell migration (Karagiannis *et al*, 2012; Gutiérrez *et al*, 2017). Additionally, expression of classic markers of epithelial cells – *CDH1*, *EPCAM*, *VIL1*, and *KRT19* are decreased in patient tumors post-treatment with radio-chemotherapy, and markers of mesenchymal cells are increased (Figure 4C). These data strongly suggest that radiochemotherapy of rectal tumors suppresses Wnt signaling and induces a more invasive, EMT-like phenotype in the surviving, wounded tissue.

**Figure 4.**
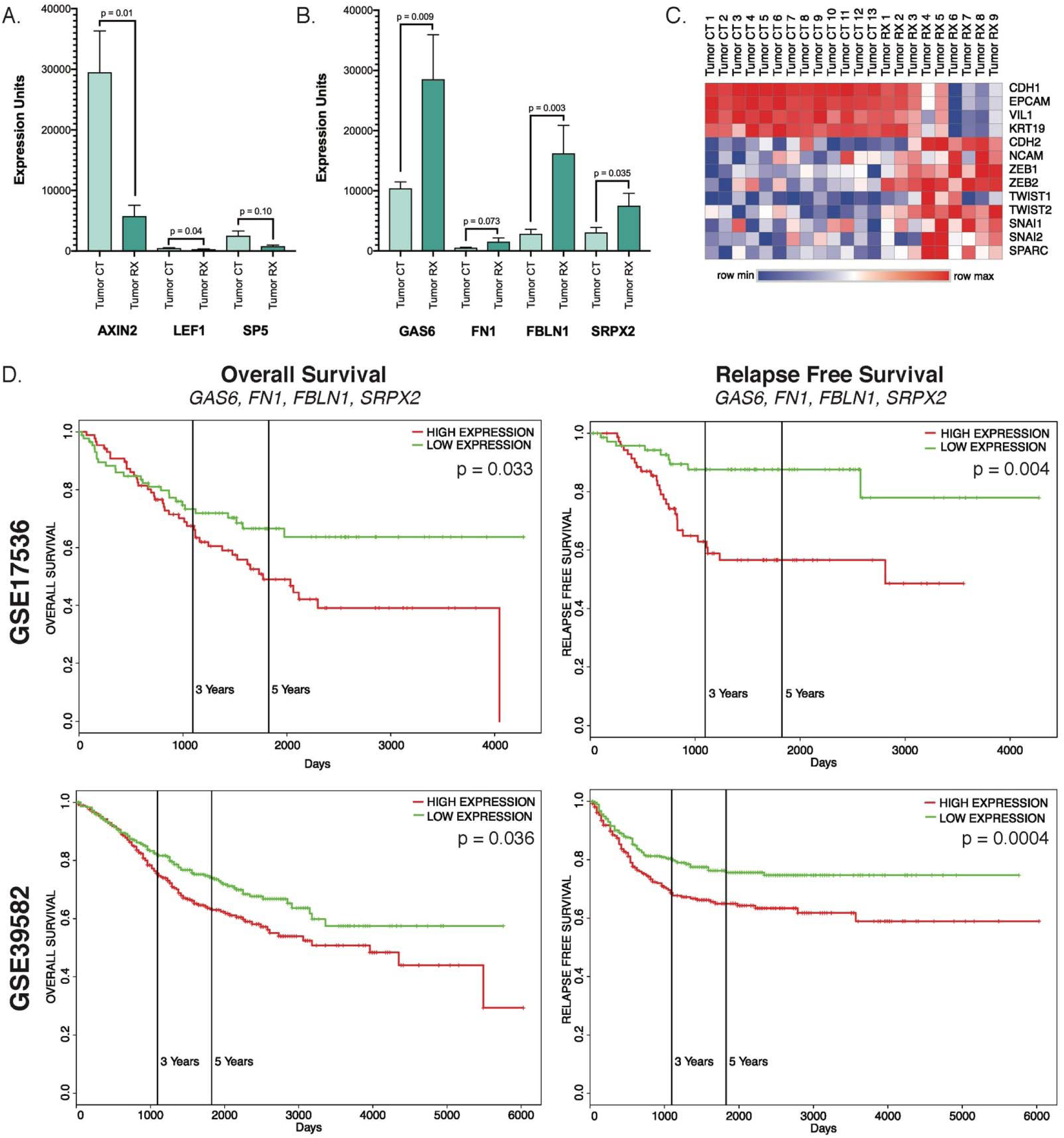
Colorectal cancer patients with elevated stromal gene expression have poorer outcomes. A. Rectal cancer patients treated with radiochemotherapy (GDS3756) display significant decrease in Wnt target gene expression compared to untreated patients. B. Rectal cancer patients treated with radiochemotherapy have significant increases in expression of *GAS6*, *FN1*, *FBLN1*, and *SRPX2*. C. Heatmap of genes associated with epithelial or mesenchymal phenotypes show a distinct transition to a mesenchymal phenotype in rectal cancer patients treated with radiochemotherapy. D. Colon cancer patients with elevated expression of *GAS6*, *FN1*, *FBLN1*, and *SRPX2* have significantly poorer overall outcome and relapse-free survival. Kaplan-Meier curves from two CRC studies (details in Supplemental Figure 7) show significant decreases in overall survival and relapse-free survival in patients overexpressing the listed genes.

Expanding the analysis to larger colorectal patient datasets, we find that elevated levels of *GAS6*, *FN1*, *FBLN1*, and *SRPX2* are correlated with significantly worse overall survival and relapse free survival in GSE17536 (N = 177) and GSE39582 (N = 586) (Study details in Supplemental Figure 7). Notably, the difference in relapse free survival curves is more significant in both studies, suggesting that elevated expression of these genes encourages disease progression to an invasive state. These genes are particularly interesting in that they have previously been connected to metastasis, but to date, they have not been linked to Wnt signaling, much less a decrease in Wnt signaling. Of the genes that we have identified, a number of them have small molecule inhibitors, though only *AXL* has inhibitors in clinical trials. None of these inhibitors have been characterized for use in colon cancer, presenting a potential new avenue for investigation.

## Discussion

Here we report that interfering with ß-catenin-dependent Wnt signaling in colon cancer cells increases their cell migration and invasion activities both *in vitro* and *in vivo*. We demonstrate this multiple ways by interfering with Wnt signal transduction at steps that lie either upstream or downstream of ß-catenin. Wnt-signaling-inhibited cell lines show increased cell migration *in vitro*, a phenotype that correlates well with *in vivo* phenotypes of increased invasiveness, including the formation of extracolonic tumors. We predict that increased invasiveness would eventually manifest as metastases were we able to carry out the experiments longer. However, the genetically manipulated cells are so invasive and aggressive at the primary site of injection in the caecum that mice became moribund and therefore limit the timeline of the study. We compared significantly upregulated genes in each of our lines to identify genes that are commonly upregulated when Wnt signaling is decreased, and discovered that a set of highly upregulated genes predicts poor outcomes in human CRC patients. We suggest that therapies targeting one or more of these genes may find clinical application in colon cancer patients presenting at the early stages of advanced disease.

Activating mutations to the Wnt signaling pathway have long been characterized as a hallmark of colon cancer, and multiple studies have pointed to the contribution of overactive Wnt signaling to metastatic disease. In patient samples, the leading edge of colon tumors stain strongly for nuclear ß-catenin, a marker of active Wnt signaling (Jung *et al*, 2001). Additionally, Snail and Twist, two genes that play key roles in EMT are direct Wnt target genes (ten Berge *et al*, 2008). Many previous studies have shown that decreasing Wnt signaling in colon cancer by targeting varying components of the pathway, can significantly reduce tumor burden in subcutaneous xenograft mouse models, the most common mouse model for pre-clinical testing of cancer therapeutics (Morin *et al*, 1996; Satoh *et al*, 2000; Tetsu & McCormick, 1999; Van de Wetering *et al*, 2002; Polakis, 2012). Indeed, studies by our group using the dnLEF1 construct has shown the same result in subcutaneous xenografts (Fig.1A-C), (Pate *et al*, 2014; Lee *et al*, 2017). Because of this, many efforts and deep resources have been brought to bear on the goal of bringing small molecule inhibitors of Wnt signaling into clinical practice. However, so far, none of these compounds have been able to show measurable benefit in colon cancer treatment, and the trials of Wnt inhibitors have shifted their focus to other cancer types (Kahn, 2014). Recent evidence has shown that the genetic landscape of colon cancer is significantly more complex than previously appreciated; a meta-analysis of patient samples found that tumors with the lowest levels of Wnt signaling had the highest levels of stromal infiltration and the worst overall patient survival (Guinney *et al*, 2015). By contrast, tumors with the highest levels of Wnt signaling had the best overall patient survival. The data that we present here supports this notion, and suggests that decreasing Wnt signaling directly favors an invasive phenotype in colon cancer.

Using gene ontology, we observe that dnLEF1 expression is associated with enrichment for genes associated with epithelial to mesenchymal transition and loss of cell adhesion. We find that high-level expression of these genes is significantly correlated with poorer patient outcomes in multiple patient data sets (Figure 4). The genes identified have been characterized to promote cell migration, either through changing gene expression or creating a more permissive extracellular matrix. Interestingly, RNAseq analysis of dnLEF1 expression in SW480 and SW620 cells cultured *in vitro* also showed changes in gene programs for cell adhesion, mobility, cell junction, and extracellular matrix (Supplemental Figure 8). However, the specific genes connected to these genes programs were different, showing that while the gene programs and cell behaviors that link to low Wnt signaling to invasive phenotypes is the same, the specific genes that change expression are different *in vitro* versus the orthotopic setting. Collectively, these programs highlight attempts by cancer cells to alter their microenvironment to promote invasion, as fibrotic, stiffened extracellular matrix has been shown to enhance EMT and invasion in many cancers (Kai *et al*, 2019).

Our findings suggest that Wnt signaling may indirectly repress invasive genes, fulfilling a specific role for Wnt signaling as an inhibitor of localized invasion in colon cancer. It is therefore possible that lower levels of Wnt signaling in APC-mutant colorectal cancer is linked to a colon cancer invasion-metastasis cascade and development of an invasion-promoting tumor microenvironment. Our results are concurrent with the CMS/CRIS colon cancer characterization studies and highlight how a decrease in Wnt signaling is an important contributing factor to advanced colon cancer and poor patient outcomes.

The ramifications of our findings are that while targeting Wnt signaling in colon cancer may reduce tumor burden, a potential, inadvertent side effect might be to induce surviving cancer cells to become invasive. Thus, as clinical trials continue to test the efficacy of newer, more specific Wnt inhibitors on Wnt-driven cancers, we suggest that treatment of patients with Wnt inhibitors should include concurrent treatment with drugs targeting one or more of the genes identified in this study, such as the GAS6:AXL pathway. Small molecule inhibitors have already been developed to target AXL and these have advanced to clinical trials. However, while AXL has been observed to be significantly overexpressed in malignant cells, none of the clinical trials currently testing AXL inhibitors are focused on colon cancer (Axelrod & Pienta, 2014). We suggest that AXL inhibitors may be therapeutically useful in concert with other drug therapies, particularly Wnt or VEGF. Small molecule approaches targeting extracellular matrix proteins and related cell-ECM interactions have been found to have mixed clinical outcomes (Kai *et al*, 2019) – given our finding that such genes are upregulated in Wnt-low colon cancer, these small molecules may be prime candidates for combination with Wnt inhibitors.

What remains unanswered from our study is the apparent “ß-catenin paradox” that our group and others have observed (Brabletz *et al*, 2001; Jung *et al*, 2001; Brabletz *et al*, 2005); that is, the heterogeneity of nuclear ß-catenin in a colon tumor, and the high expression at the invading edge of colon cancer. We speculate that this may be due to a temporal function of tumor invasion; as high Wnt signaling promotes growth and proliferation, the decrease in Wnt signaling to promote invasion may be a transient state.

Progression to metastatic disease remains the most challenging aspect of colon cancer treatment, and the one in which patient outcomes remain poor overall. The finding that these advanced tumors harbor relatively lower levels of Wnt signaling compared to other colon cancer subtypes, points to a previously uncharacterized role of Wnt signaling in colon cancer. Our study demonstrates that decreasing Wnt signaling induces a more invasive phenotype, and identifies a number of contributors to this invasive state. We suggest that future studies of Wnt inhibitors for clinical use consider concurrent treatment with inhibitors of these identified targets, to both reduce tumor burden and prevent cancer invasion.

## Materials and Methods

### Cell lines

SW480, SW620, COLO320, and HCT116 were obtained from ATCC. Cell lines were cultured in Dulbecco’s Modification of Eagle’s Medium (DMEM, Hyclone) or RPMI-1640 (Hyclone) supplemented with 10% Fetal Bovine Serum (Atlas Biologicals), 1% Penicillin/Streptomycin (Mediatech), and 2mM Glutamine (Mediatech). Unless otherwise stated, all assays were conducted in the same media. dnLEF1 cell lines were prepared as previously described (Pate *et al*, 2014). LRP6 knockout (LRP6KO) cell lines were created by co-transfecting cell lines with 3 μg LRP6 CRISPR and 300 ng eGFP-Puro using BioT transfection reagent (Bioland Scientific, B01). After selection with Puromycin, cells were sorted for LRP6 knockdown using flow cytometry (anti-human LRP6-APC, R&D Systems MAB1505) and successful knockdown was confirmed by Western blot.

### LRP5 and LRP6 CRISPR constructs

The CRISPR kit used for constructing multiplex CRISPR/Cas9 vectors was a gift from Takashi Yamamoto (Addgene kit #1000000054). Guide RNAs targeting LRP5 and LRP6 were designed using the Zhang Lab Optimized CRISPR Design Tool (http://crispr.mit.edu). A Cas9 nickase system was used such that two closely aligned guides per gene were developed.

For LRP5, the primer pairs used were:

LRP5-Cas9n-1-Fwd: CACCGCTCGGTCCAGTAGATGTAGC

LRP5-Cas9n-1-Rev: AAACGCTACATCTACTGGACCGAGC

LRP5-Cas9n-2-Fwd: CACCGCGGCAAGCCGAGGATCGTGC

LRP6-Cas9n-2-Rev: AAACGCACGATCCTCGGCTTGCCGC.

For LRP6, the primer pairs used were:

LRP6-Cas9n-1-Fwd: CACCGTTGGCCAAATGGTTTAGCCT

LRP6-Cas9n-1-Rev: AAACAGGCTAAACCATTTGGCCAAC

LRP6-Cas9n-2-Fwd: CACCGAAGTGTTAACCAATACTACA

LRP6-Cas9n-2-Rev: AAACTGTAGTATTGGTTAACACTTC.

Guides were inserted into vectors using T4 polynucleotide kinase (New England Biolabs), and the final combined guide RNA vector was assembled by Golden Gate Assembly using Quick Ligase (New England Biolabs). The final vector was electroporated into DH5α competent cells and purified using a Nucleospin Maxiprep kit (Macherey Nagel). Restriction enzyme digested products were run out on an agarose gel to confirm insert size and plasmid quality. Correct assembly was verified by Sanger Sequencing.

### LRP5 and LRP6 shRNA

LRP5 & LRP6 shRNA constructs were obtained from the RNAi Consortium database (Sigma Aldrich). Lentivirus was packaged in HEK293T cells (ATCC) and purified using PEG-it (System Biosciences).

### Western blotting

Whole cell lysates were prepared by resuspending a cell pellet in an appropriate amount of RIPA buffer containing protease inhibitors and phosphatase inhibitors and incubating for 30 minutes on ice. The lysates were spun down at high speed for 15 minutes at 4C, and the supernatant was transferred to a clean tube. Lysates were quantified using a Bradford assay (BioRad, 500-0205). 80 μg of total cell lysates were analyzed by Western blot using the following antibodies and concentrations: LRP6 (1:1000, Cell Signaling Technology 3395), LRP5 (1:1000, Cell Signaling Technology 5731), ß-Catenin (1:1000, Cell Signaling Technology 8480), ß-Tubulin (1:2000, Genetex GTX101279). All blots were incubated overnight in primary antibody, washed, and then incubated for two hours in secondary antibody – anti-rabbit-HRP (1:5000, GE Healthcare) or anti-mouse-HRP (1:2000, GE Healthcare). Blots were imaged using a Syngene G-Box system. Bands were quantified using Adobe Photoshop. Statistical evaluation of three or more independent biological replicates was performed using Student’s unpaired T-Test.

### Luciferase assay

Cells were seeded at a density of 1 *10^5^ cells per well in a 12-well tissue culture plate using antibiotic-free media 24 hours prior to transfection. Cells were co-transfected with 100ng Wnt ligand expression plasmid (Najdi *et al*, 2012), 300ng LRP5 or LRP6 CRISPR construct, 100ng SuperTOPFlash or SuperFOPFlash (Gifts from Dr. R.T. Moon, Addgene plasmid 12456) and 100ng CMV-ß-Galactosidase using BioT (Bioland Scientific B01-02) per manufacturer’s instructions. Cells were harvested 24 hours post-transfection and assayed for luciferase activity and ß-galactosidase activity (as a normalization control). Statistical evaluation was performed using Student’s unpaired T-test.

### Scratch assay

Cells were seeded at a density of 2*10^6^ cells per well in a 6-well plate. After 24 hours, 2 crosses were scratched into the cell monolayer with a P1000 tip. Each well was washed once with PBS before incubating in media. Images were taken at 0 hours, 24 hours, and 48 hours post-scratch. Measurements were obtained using Adobe Photoshop image analysis. Statistical evaluation was performed using Student’s unpaired T-test.

### Flow cytometry

One million cells per sample condition were collected and washed with FACS buffer (3% FBS in PBS). If needed, cells were fixed in 4% paraformaldehyde for 15 minutes at room temperature, permeabilized by 0.1% saponin in HBSS for 15 minutes on ice, and washed twice with FACS buffer prior to staining. Cells were incubated with primary antibody for 1 hour at room temperature in the dark. If secondary antibody was needed, cells were washed twice with FACS buffer prior to incubating in secondary antibody for 30 minutes at room temperature, in the dark. Cells were finally washed and resuspended in 300-600 μL FACS buffer per sample and analyzed on either a BD FACSAria or Acea Biosciences Novocyte. Data was analyzed using FlowJo (Treestar).

### Orthotopic injections

Immuno-deficient NSG mice (Jackson labs) were anaesthetized with 300μl of 100mg/kg ketamine/10mg/kg xylazine. The surgical site was shaved and cleaned with ethanol. After an incision was made, the caecum was drawn out and washed with sterile PBS. Cells (5*10^5^) were injected in the caecum wall at six to eight sites using a Hamilton syringe (Hamilton 80301), three to four injections per side. The peritoneal wall and skin were sutured separately, the former with resorbable sutures, the latter with nylon. 5mg/kg carprofen was injected as an analgesic. Mice health was monitored for 28 days prior to harvest.

### Mesentery dissociation for flow cytometry

Mesentery from orthotopically-injected mice were collected in a petri dish on ice and briefly diced with razor blades before transferring the tissue pieces into a 40 micron strainer and pushing tissue through the strainer using the plunger of a 3ml syringe. The strainer was rinsed with 10ml DMEM with 5% FBS, 1% P/S, and 2mg/ml Collagenase I (Sigma). The mixture was collected in a conical tube and shaken at 37 C for 1 hour. The tube was spun down and the supernatant aspirated. The pellet was washed in 10ml HBSS, spun down, and supernatant aspirated except for approximately 500 μl of HBSS. 50 μl DNAse I (10 U, Thermo Fisher) was added and the pellet resuspended and incubated at room temperature for five minutes. Following DNAse treatment, 2 ml of 0.05% trypsin was added, and the mixture was incubated at 37 C for 10 minutes. 5 ml DMEM with 5% FBS was then added, the mixture spun down, and the supernatant was aspirated. The pellet was resuspended in 5 ml DMEM and passed through a 40-micron cell strainer. The strainer was washed with an additional 5 ml of DMEM and the cell number and viability was measured using a Countess II (Thermo Fisher Scientific). 1*10^6^ cells per sample were incubated in CD298-APC (Machery Nagel) for one hour at room temperature in the dark. Samples were washed twice in FACS buffer before resuspending in FACS buffer and running on an Acea Biosciences Novocyte. Data was analyzed using FlowJo (Treestar).

### Subcutaneous injections

Immuno-deficient NSG mice (Jackson labs) were anaesthetized with 300μl of 100mg/kg ketamine/10mg/kg xylazine. The surgical site was shaved and cleaned with ethanol. 2*10^6^ cells were resuspended in 100 μL PBS and injected under the skin on both flanks of the mouse. Mice health was monitored for 21 days prior to harvest.

### Immunohistochemistry

Excised caecums were fixed in 10% formalin for 24 hours, cut on the latitude, and mounted on edge in paraffin. 10μm FFPE sections were cut onto SuperFrost Plus slides. For antigen retrieval, slides were deparaffinized and rehydrated, followed by antigen retrieval in a pressure cooker using 10mM sodium citrate for five minutes at pressure. Slides were then stained by hematoxylin and eosin, dehydrated and mounted using Permount mounting medium (Fisher). Slide images were captured with a Keyence BZ-X700 system and processed in Adobe Photoshop.

### Pathology scoring

A blinded evaluation of stained slides was performed by pathologists who scored each section of caecum on the absence, low presence, or high presence of phenotypic features. Additional measurements were taken of each section by comparing the total pixels of a section to pixels of individual components (eg. extracolonic tumors, intact epithelia, et cetera) and normalizing by percentage of total section. Approximately three to five sections per caecum from at least three mice per cell line were analyzed. The complete set of analyzed images is in Supplemental Materials Figure 1.

### Total RNA Isolation

RNA was extracted from cells using TRIzol (Invitrogen) and DirectZol RNA Extraction Kit following the manufacturer’s instructions (Zymo Research). RNA was extracted from flash-frozen tissue samples by using a mortar and pestle to crush tissue into fine powder, homogenizing in TRIzol using a Precellys 24 (Bertin), and extracting the RNA with the DirectZol kit. RNA quality was checked with Agilent Bioanalyzer. Samples with RNA Integrity Number (RIN) scores ≥ 9 were used for RNA-seq library construction.

### Quantitative PCR

RNA was extracted from cells using TRIzol (Invitrogen) and DirectZol RNA Extraction Kit following the manufacturer’s instructions (Zymo Research). RNA was extracted from flash-frozen tissue samples by using a mortar and pestle to crush tissue into fine powder, homogenizing in TRIzol using a Precellys 24 (Bertin), and extracting the RNA with the DirectZol kit. cDNA was synthesized from 1 μg of total RNA using the High Capacity cDNA Reverse Transcription Kit (Invitrogen), as per the manufacturer’s instructions. qPCR was performed in triplicate for each experimental condition using Maxima SYBR Green qPCR Master Mix (Invitrogen), according to the manufacturer’s instructions. Primer pairs used are as follows:

*AXIN*: Forward: CTGGCTTTGGTGAACTGTTG Reverse: AGTTGCTCACAGCCAAGACA

*MYC*: Forward: CTACCCTCTCAACGACAGCA Reverse: AGAGCAGAGAATCCGAGGAC

*SP5*: Forward: AATGCTGCTGAACTGAATAGA Reverse: AACCGGTCCTAGCGAAAACC

*GAPDH*: Forward: TCGACAGTCAGCCGCATCTTCTT Reverse: GCGCCCAATACGACCAAATCC.

### Library Construction

RNA-seq libraries were built using the Smart-seq2 protocol (Picelli *et al*, 2014) with modifications according to (Serra *et al*, 2018). Briefly, for each sample 10 ng of total RNA was converted to full-length cDNA using poly-dT primer and reverse transcriptase. The full-length cDNA was amplified using nine PCR cycles. 18 ng full-length cDNA for each sample was converted to sequencing library by tagmentation using the Illumina Nextera kit. Eight PCR cycles were used for library amplification. The libraries were multiplexed and sequenced on an Illumina NextSeq500 sequencer as 43 bp paired-end reads.

### Analysis

Adapter sequences and low quality base pairs from the 5’ and 3’ ends of the paired-ends reads were trimmed using Trimmomatic v. 0.35 (Bolger *et al*, 2014) using the following parameters: “PE read1.fastq read2.fastq pe_read1.fastq.gz se_read1.fastq.gz pe_read2.fastq.gzse_read2.fastq.gz ILLUMINACLIP:NexteraPE-PE.fa:2:30:8:4:true LEADING:20 TRAILING:20 SLIDINGWINDOW:4:17 MINLEN:30”. For each sample, the reads from the host (mouse) and the graft (human) were classified and separated using Xenome (Conway *et al*, 2012). Xenome outputs five classes of reads: host, graft, ambiguous, both, and neither. Only the host and graft reads were kept for mapping to the host and graft transcriptomes respectively while the other three classes of reads (ambiguous, both, and neither) were discarded. We used GENCODE v. 28 reference transcriptome for human and GENCODE v. M18 for mouse (Frankish *et al*, 2018). RNA-seq reads for each sample were mapped to the reference transcriptome using Bowtie v. 1.2 (Langmead *et al*, 2009) with the following parameters: “-X 2000 -a -m 200 -S --seedlen 25 -n 2 -v 3”. Gene expression levels and read counts were obtained using RSEM v. 1.2.31 (Li & Dewey, 2011).

### Differential Expression Analysis

Differential gene expression analysis was performed using DESeq2 R package (Anders & Huber, 2010). Dataset overlap was performed using the online Venn diagram tool at http://bioinformatics.psb.ugent.be/webtools/Venn/. Gene ontology was performed using Enrichr (Chen *et al*, 2013; Lachmann *et al*, 2016). Network analysis performed using GeNets (Li *et al*, 2018).

### Patient survival analyses

Kaplan-Meier analyses of publically available colon cancer datasets were performed using PROGgeneV2 (Goswami & Nakshatri, 2014), SurvExpress (Aguirre-Gamboa *et al*, 2013), and cBioPortal (Cerami *et al*, 2012; Gao *et al*, 2013).

## Supplementary Materials

**Supplemental Figure 1.**
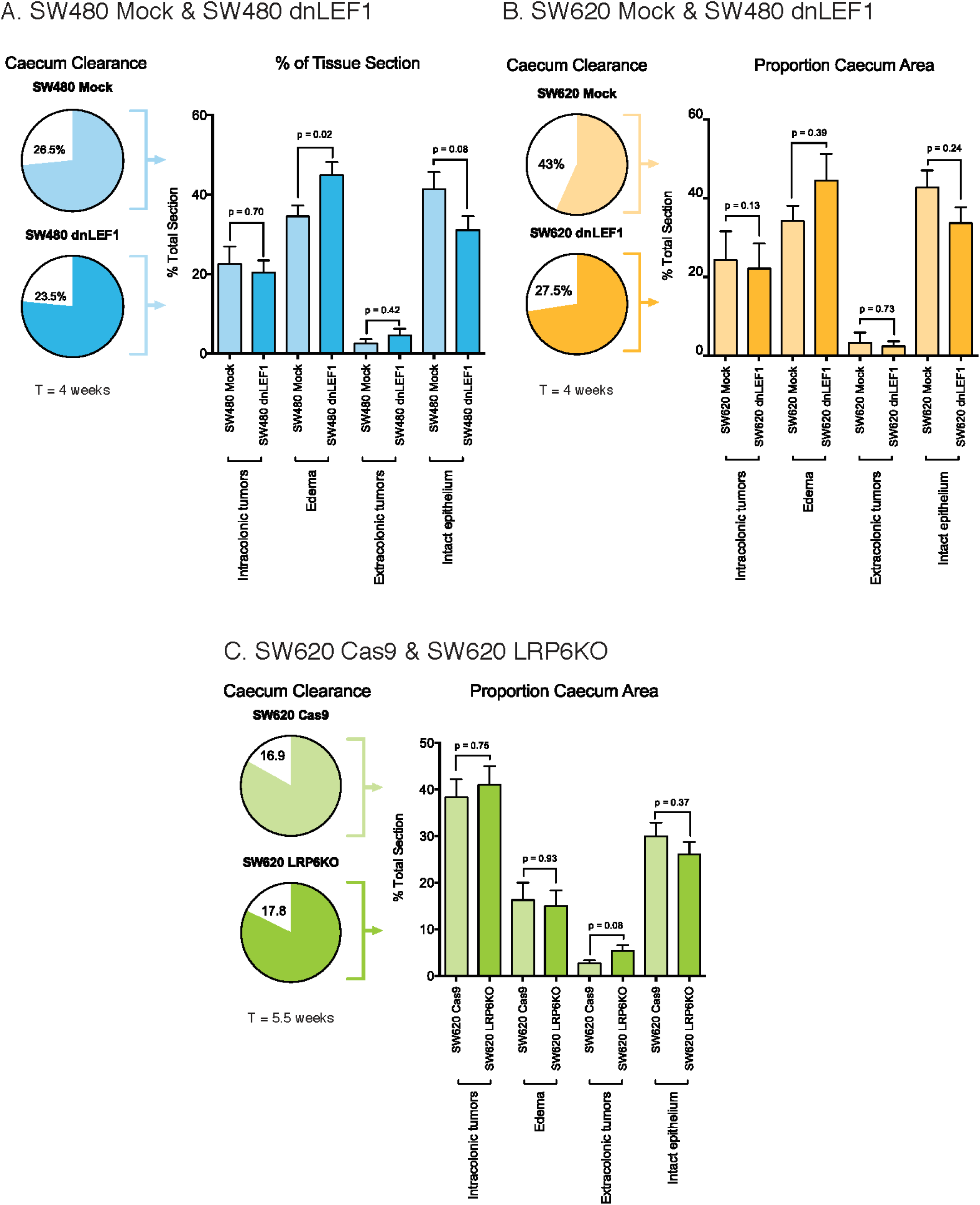
Orthotopic tumor sections for blinded pathology scoring.

**Supplemental Figure 2.**
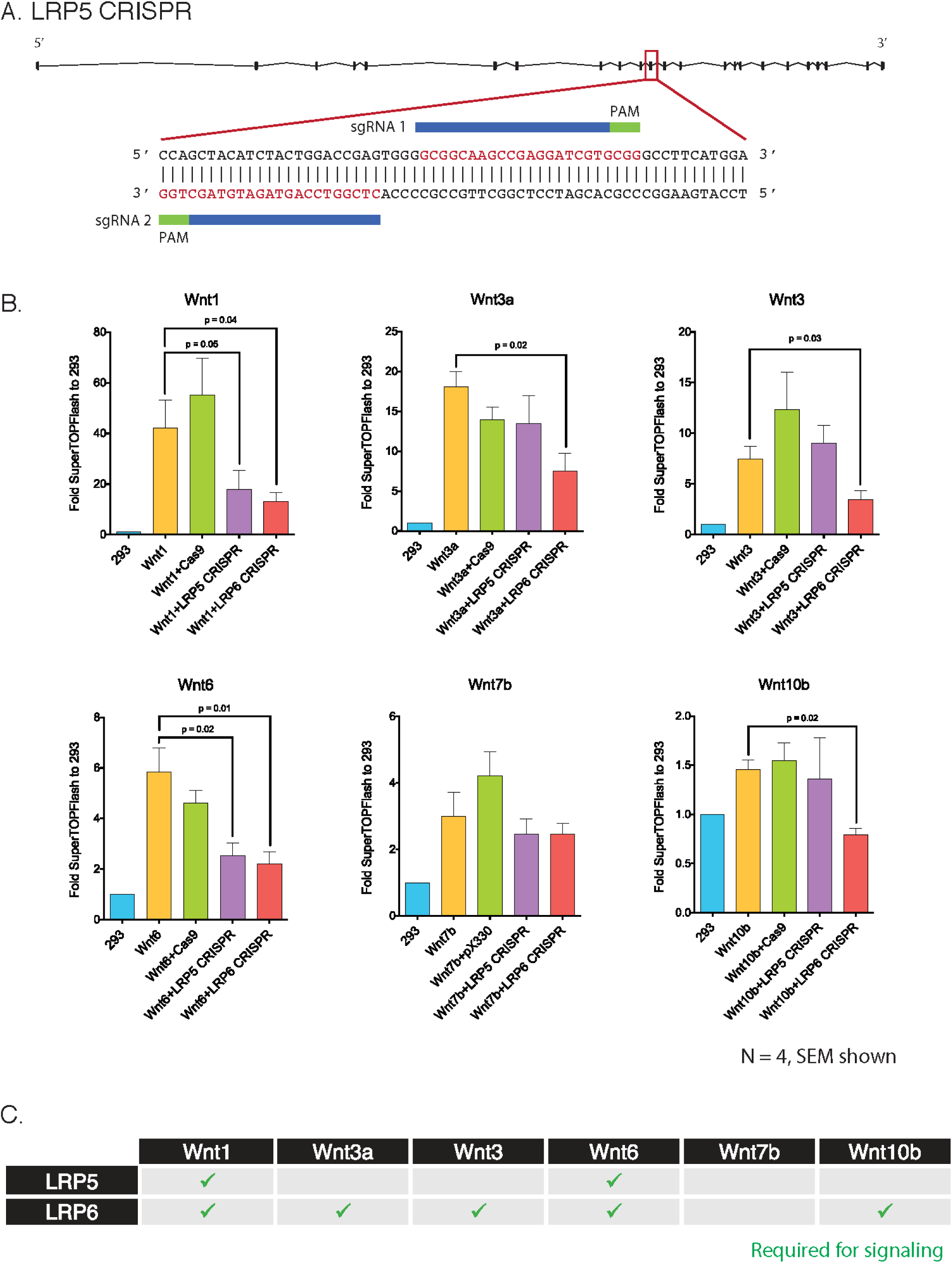
Quantification of orthotopic tumors. A. Percent of lumen clearance shows no significant difference between SW480 Mock and SW480 dnLEF1. A significant increase in edema and a decrease in intact epithelia are seen in SW480 dnLEF1. B. Percent of lumen clearance shows a significant decrease in cleared area in SW620 dnLEF1 compared to SW620 Mock. An increase in edema and a decrease of intact epithelia are seen in SW480 dnLEF1, but not to significance. C. No significant difference in lumen clearance observed in SW620 LRP6KO, but an increase in extracolonic tumors is observed.

**Supplemental Figure 3.**
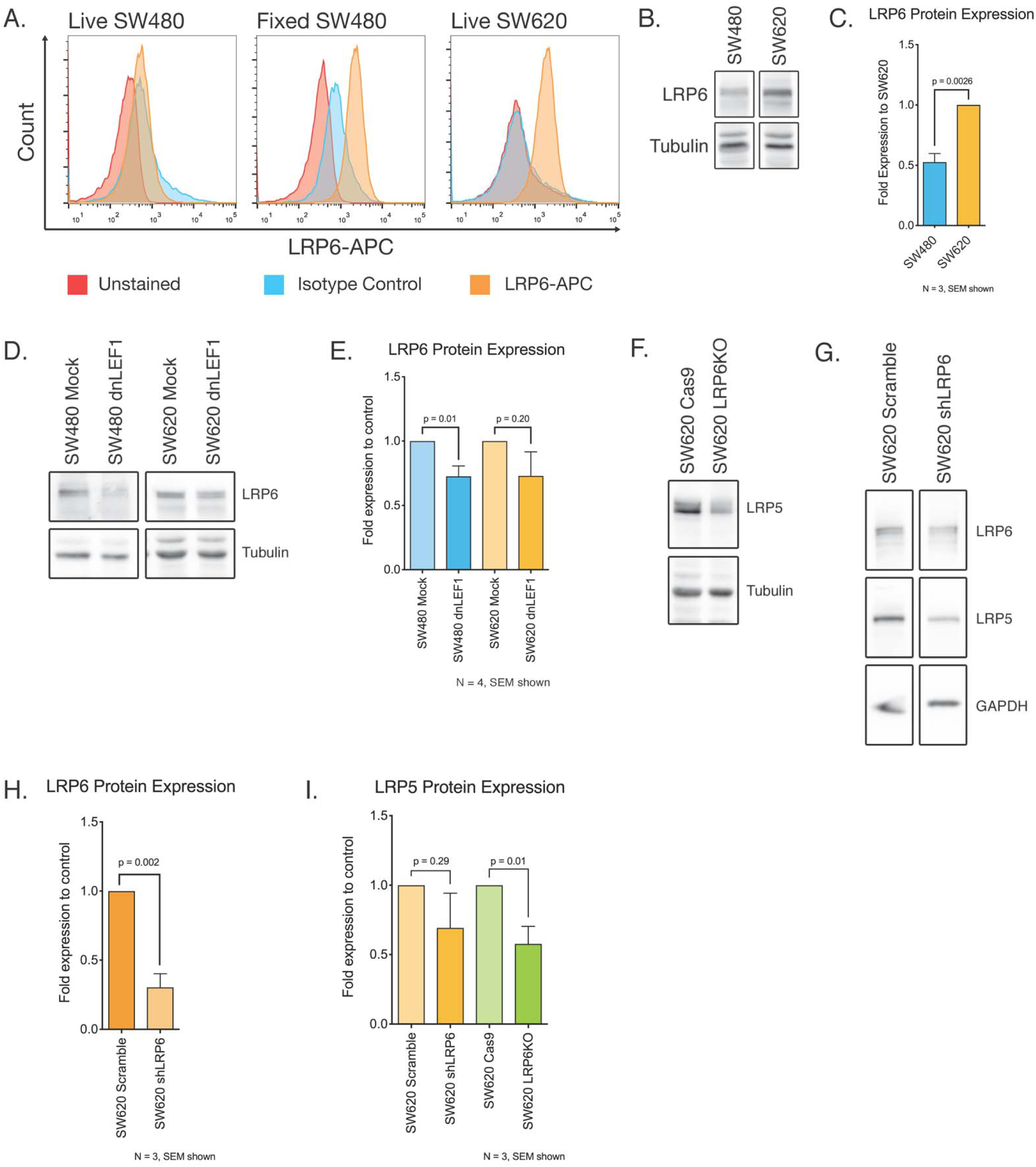
Wnt ligands require LRP6 to activate Wnt signaling. A. LRP5 CRISPR construct. B. HEK293 cells were transiently transfected with a Wnt ligand and a LRP5/6 CRISPR construct. Of the ligands tested, all except Wnt7b showed a significant decrease in Wnt signaling activity with the loss of LRP6. A number of Wnt ligands also showed sensitivity to loss of LRP5, however, no Wnt ligand was solely active through LRP5. Additionally, loss of LRP5 or LRP6 decreased Wnt activity by more than 50%, suggesting that HEK293 cells were not simply switching reliance to the other LRP co-receptor. We find that these ligands require both LRP5 and LRP6 to activate signaling. C. Summary of Wnt ligand requirements for LRP5 and LRP6.

**Supplemental Figure 4.**
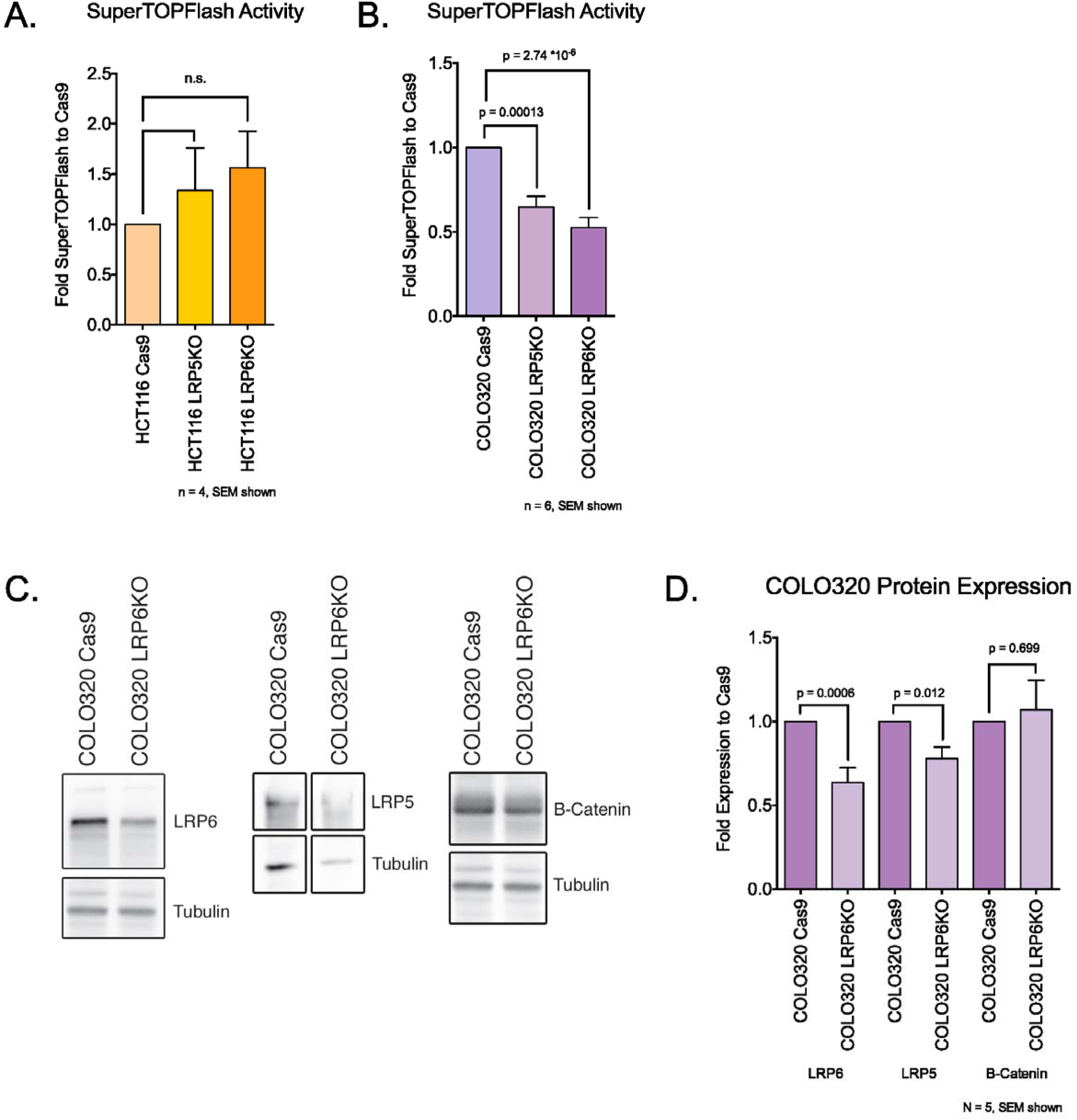
LRP6 is not expressed on the cell membrane of SW480 cells. A. Flow cytometry of SW480 and SW620 cells reveals that LRP6 is not detectable in SW480 cells, but a signal is clearly detected after cells were fixed and permeabilized, indicating LRP6 protein is present but not localized to the cell membrane. B-C. Western blot of whole cell lysates from SW480 and SW620 cells probed for LRP6 show that SW620 cells express significantly more LRP6 protein than SW480 cells. D-E. LRP6 protein expression is decreased in dnLEF1-expressing cell lines Total LRP6 protein is significantly decreased in SW480 dnLEF1 cells compared to Mock. A noticeable but not significant decrease in LRP6 is observed in SW620 dnLEF1 cells. F-H. Loss of LRP6 induces a loss of LRP5 in SW620 cells F. By Western blotting, we find a significant decrease in LRP5 protein in LRP6 knockout cells. To confirm that this was not an off-target effect of our construct, we used a lentiviral shRNA to knockout LRP6. Total LRP5 was also decreased in the shRNA-transduced cells.

**Supplemental Figure 5.**
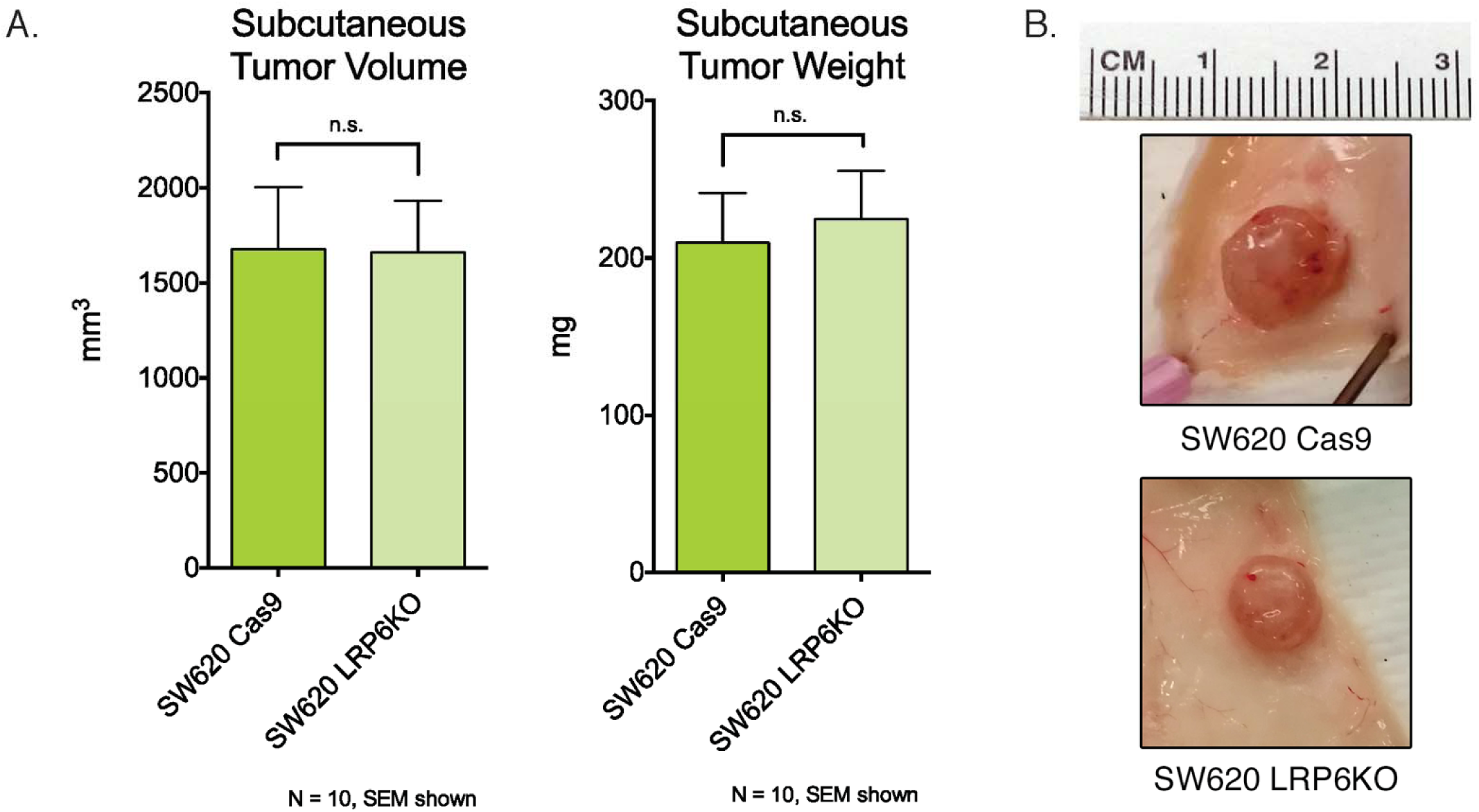
Loss of LRP6 reduces Wnt signaling in COLO320, but not HCT116 cells. A. HCT116 cells do not show a significant change to Wnt signaling activity in response to loss of LRP5 or LRP6. HCT116 cells have a mutation in b-catenin that allows the cells to evade the destruction complex in order to activate Wnt target genes. This suggests that in HCT116 cells, Wnt ligands may play a negligible role in modulating signaling activity. B. COLO320 cells show a significant decrease in Wnt signaling activity with the loss of LRP5 or LRP6. C-D. Western blots and quantitation of COLO320 Cas9 and LRP6KO cells shows a decrease in total LRP6 protein as expected. Interestingly, a loss of LRP5 is also observed with the knockout of LRP6. This is also observed in SW620 cells (See Supplemental Figure 4). Unlike the SW620 cells, we do not find a significant decrease in b-catenin in the COLO320 LRP6KO cells, suggesting a different mode of Wnt signaling regulation.

**Supplemental Figure 6.**
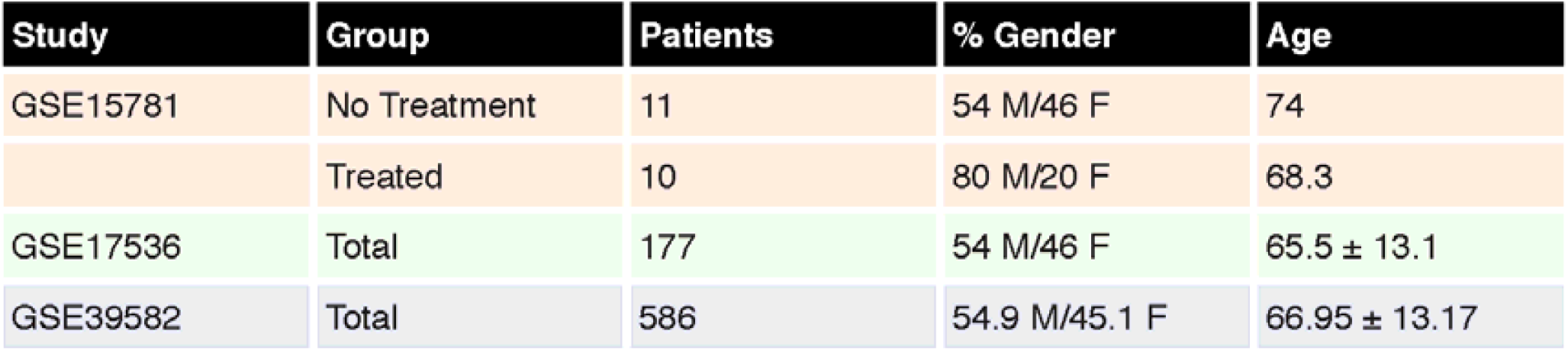
Subcutaneous tumors of SW620 Cas9 and LRP6KO. A. No significant difference in tumor volume and weight is seen between SW620 Cas9 and SW620 LRP6KO. B. Representative images of tumors at harvest.

**Supplemental Figure 7.**
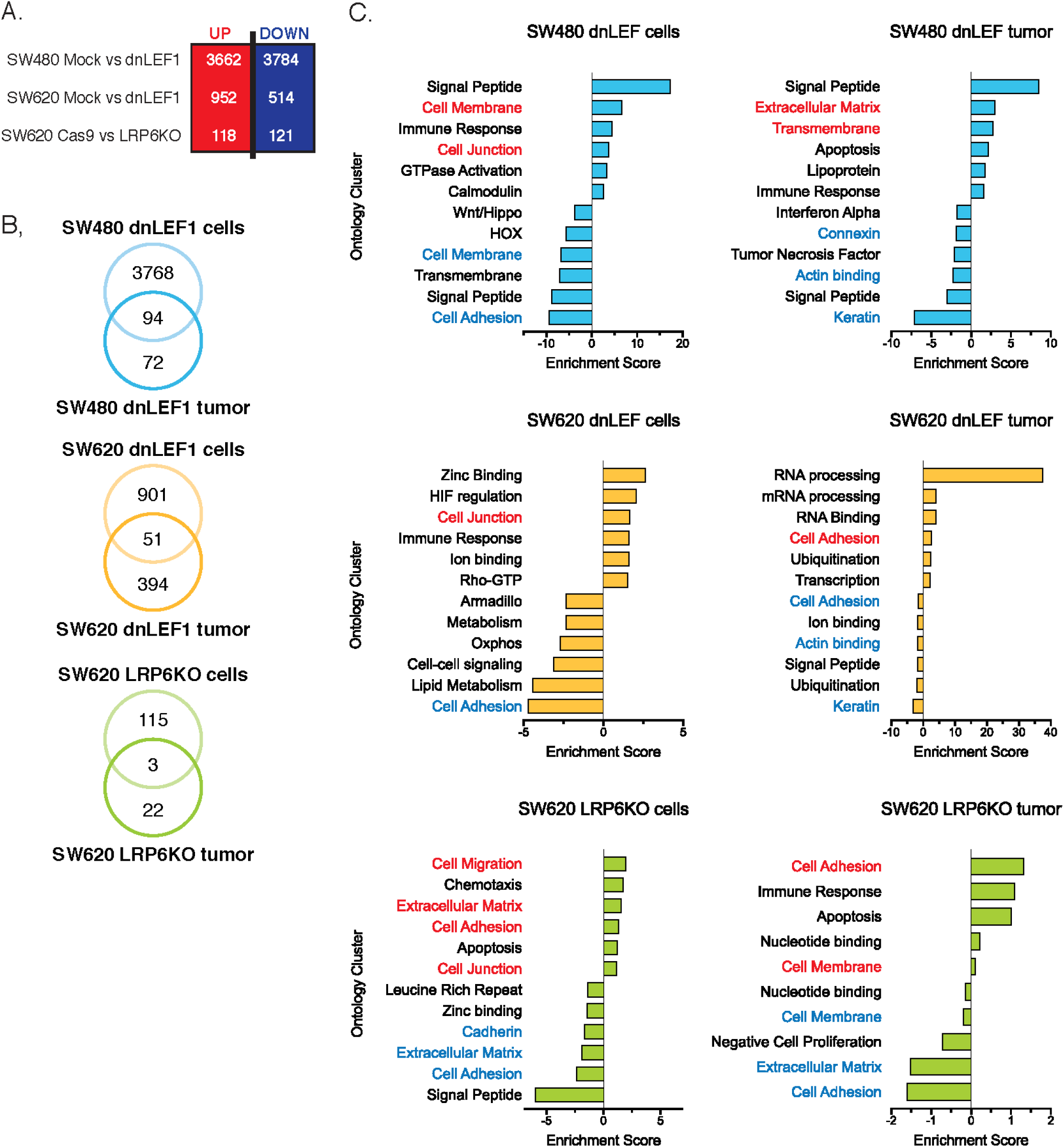
Patient characteristics from studies used in Kaplan-Meier curves. (figure 4).

**Supplemental Figure 8. Decreased Wnt signaling induces global transcriptome changes *in vitro* in genes associated with cell adhesion and extracellular matrix.** A. Significantly changed genes between Wnt-decreased cell lines and their associated control. B. Overlap of significantly upregulated genes between RNA from cell culture and RNA from orthotopic tumors shows relatively few conserved gene expression changes. C. Top DAVID gene ontology clusters for significantly upregulated and downregulated genes *in vitro* and *in vivo* reveal conserved changes in gene programs associated with cell adhesion and extracellular matrix.

Data files S1-S#

Data file S1. Significantly upregulated and downregulated genes when Wnt signaling is decreased conserved between SW480 dnLEF1 and SW620, relative to SW480 Mock.

## Acknowledgments

The authors would like to thank the members of the Waterman and Donovan labs for their scientific discussion and feedback. We would also like to thank Eric Stanbridge, Jenny Wu, Melanie Oakes, Christopher Hughes, Emiliana Borrelli, John Lowengrub, Arthur Lander, Michael McClelland, and Harry Mangalam for providing advice and critiques.

## Author contributions

GTC and MLW conceived of the experiments and wrote the manuscript. GTC and DFT performed the cell biology and animal experiments. GTC and YL created the cell lines and performed the experiments. RM and AM performed the orthotopic tumor sequencing and analysis. RAE and DFT performed the blinded pathology scoring and obtained all animal protocol approvals.

## Data and materials availability

The data discussed in this publication have been deposited in NCBI’s Gene Expression Omnibus (Edgar *et al*., 2002) and are accessible through GEO Series accession number GSE130236 (https://www.ncbi.nlm.nih.gov/geo/query/acc.cgi?acc=GSE130236).”

## Supplementary Materials for Chen *et al*

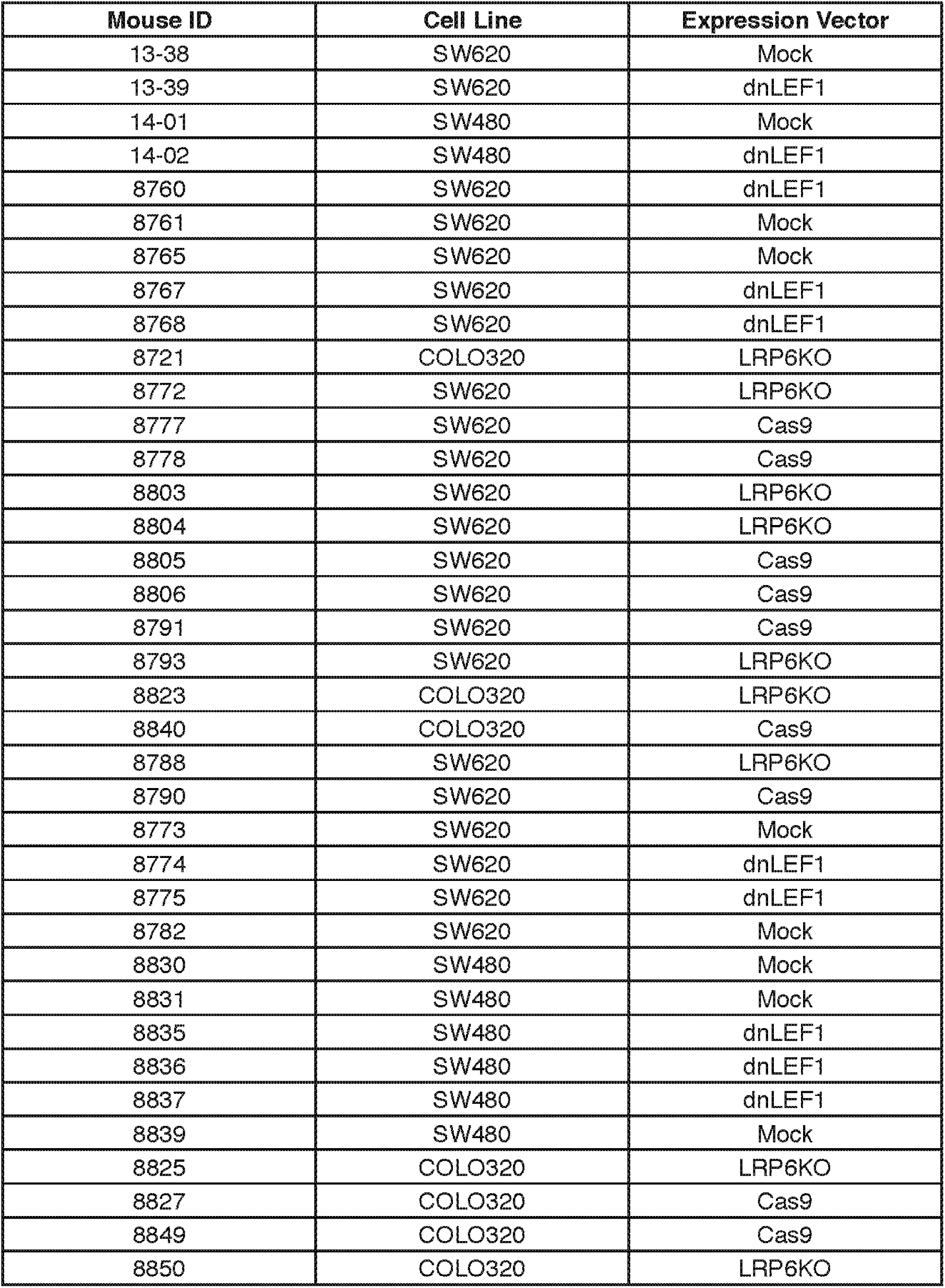
Key for orthotopic tumor sections used in pathology scoring.

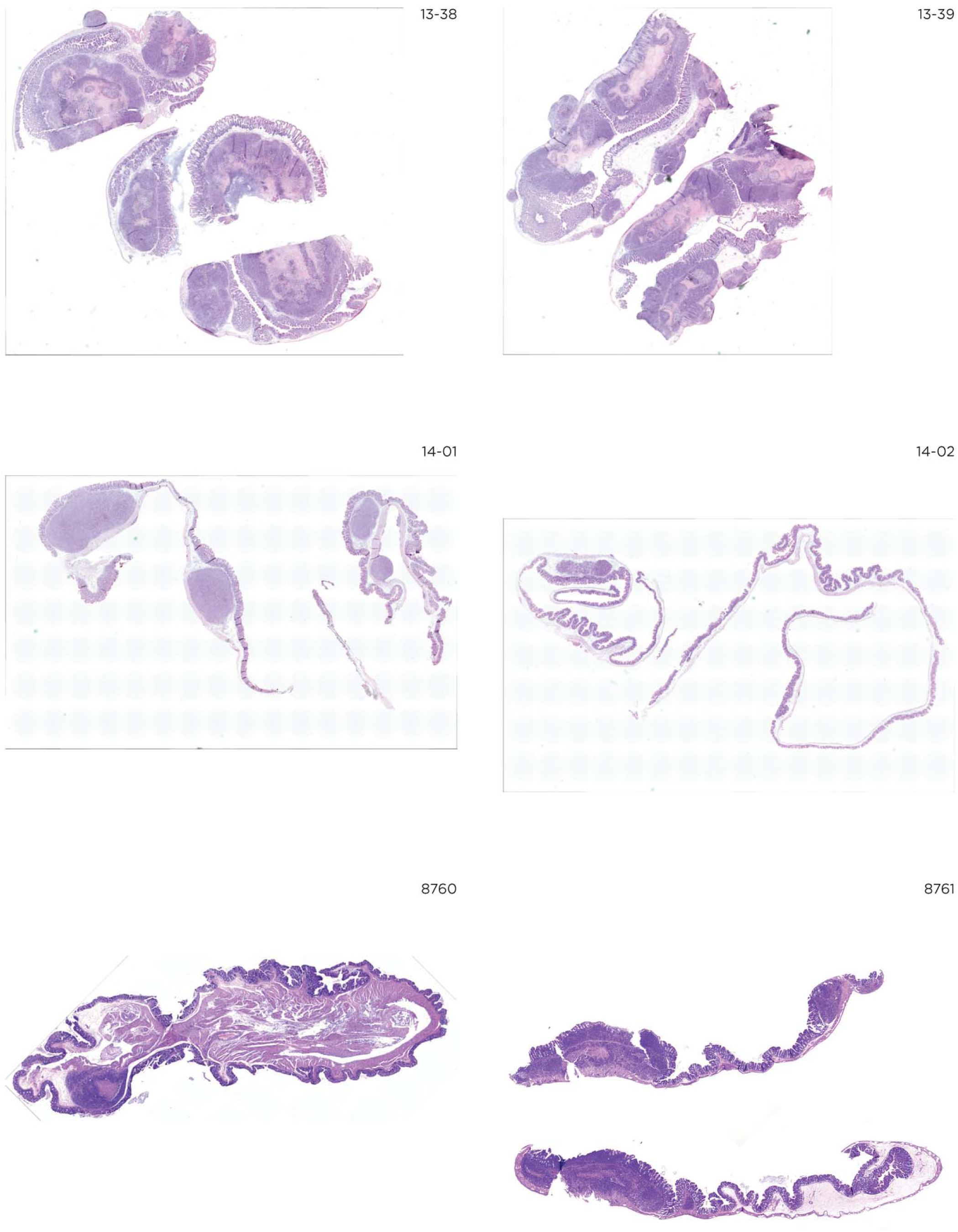

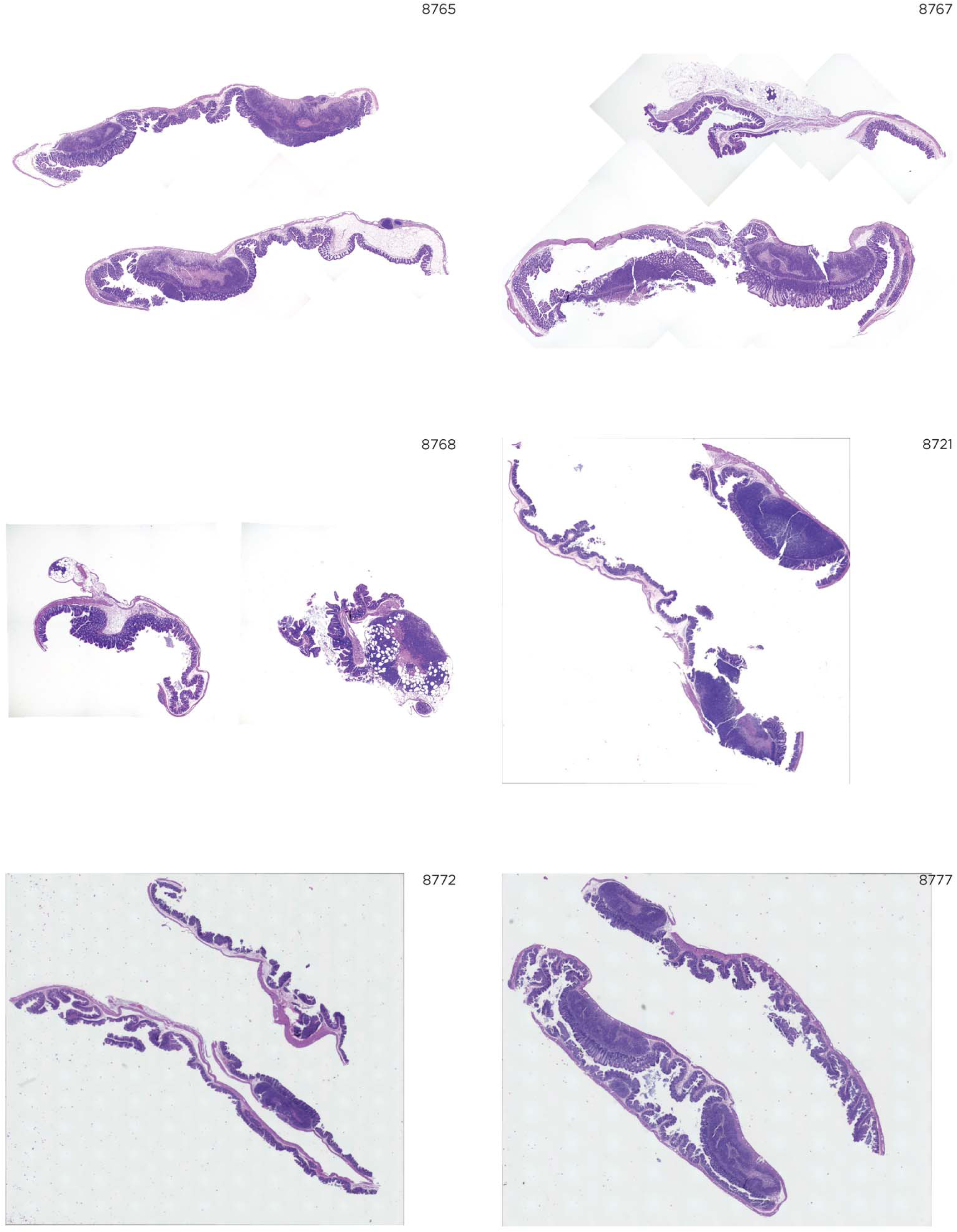

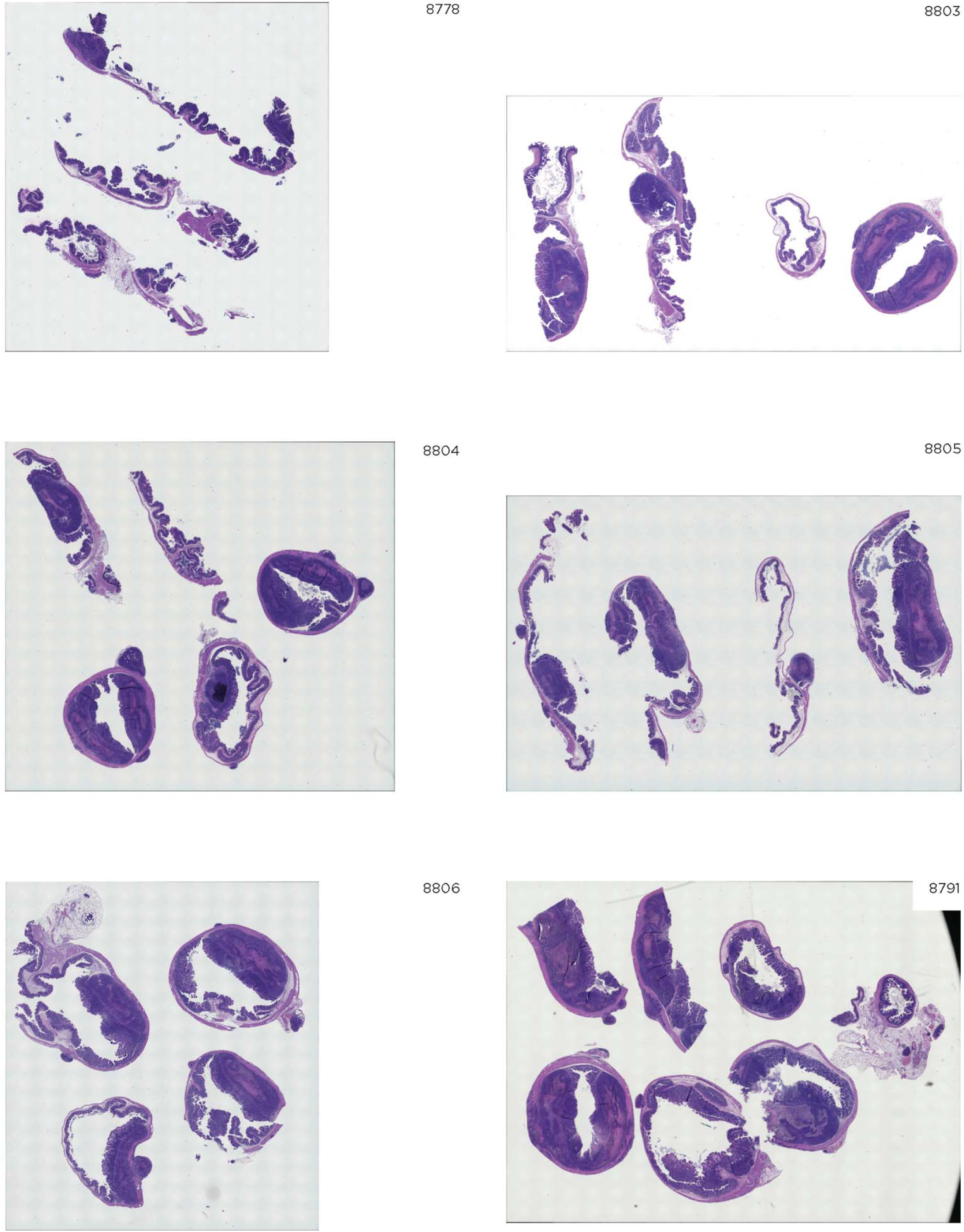

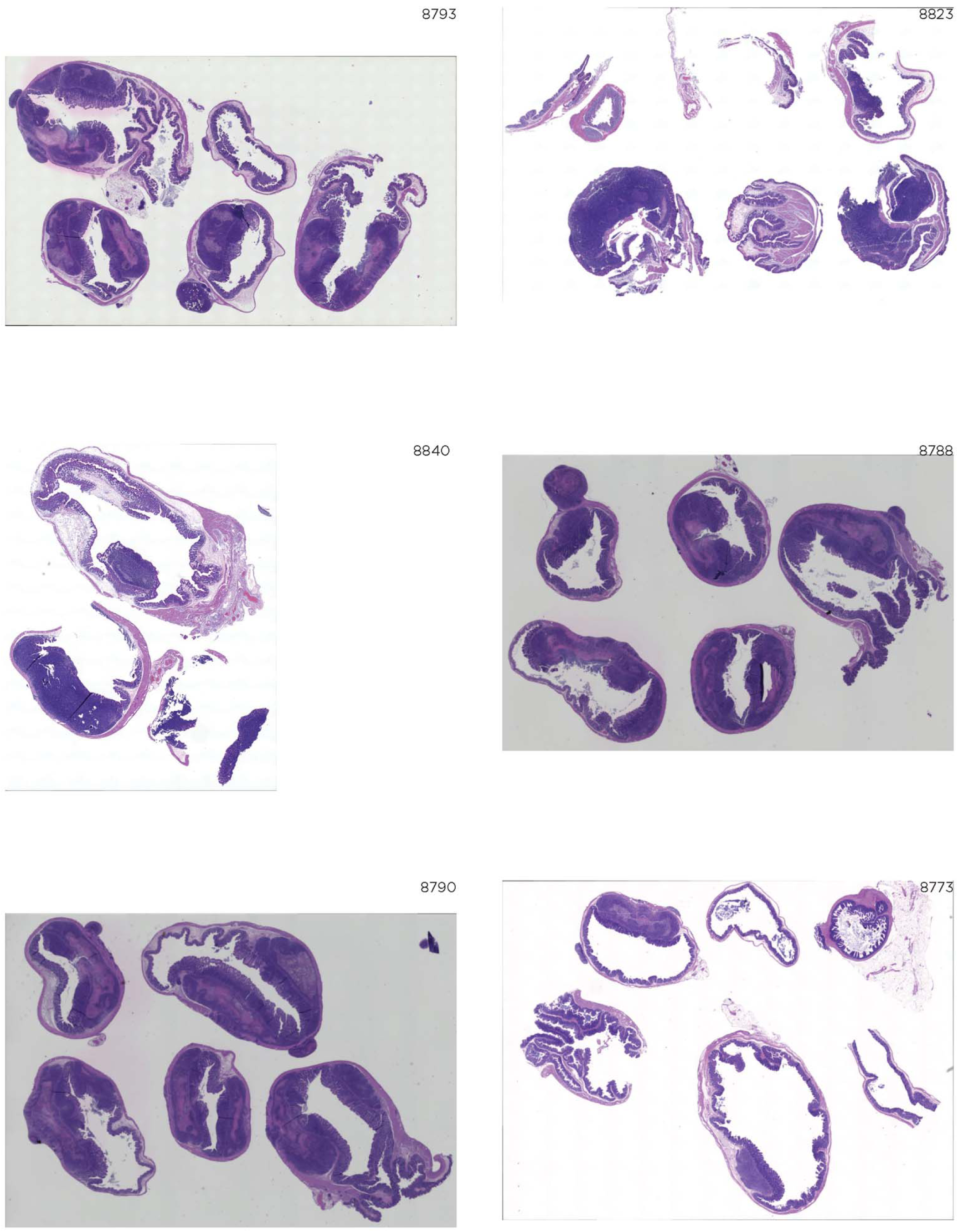

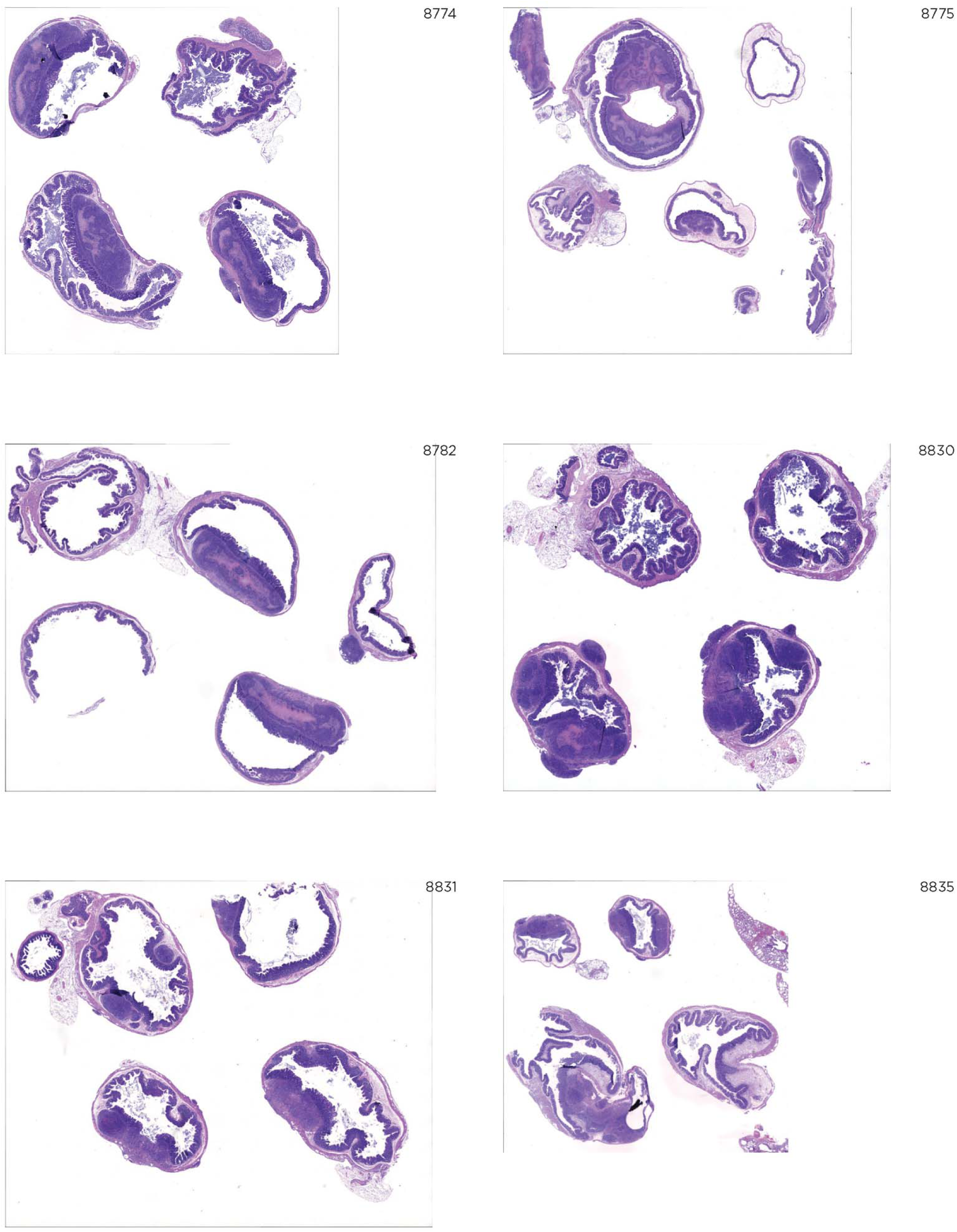

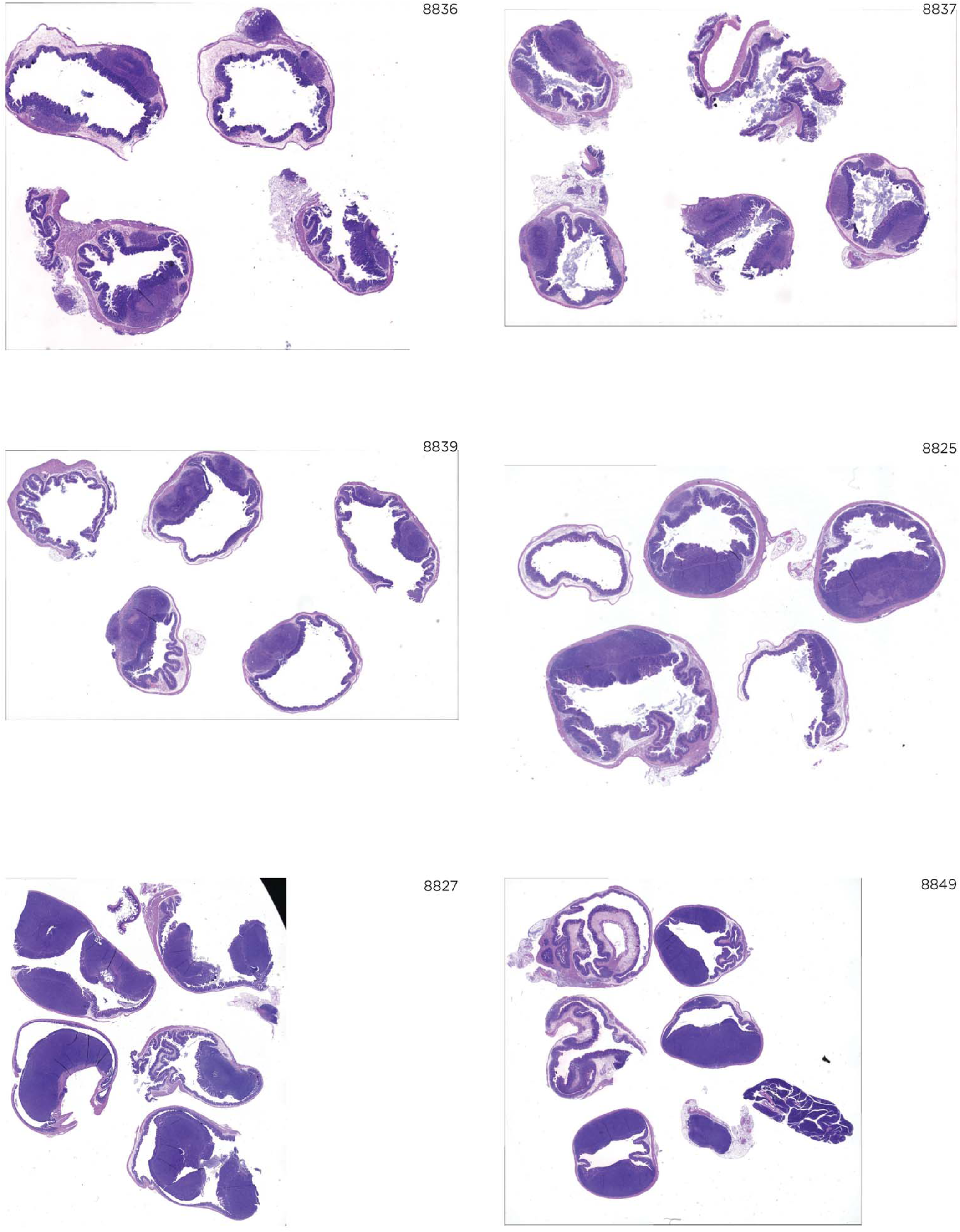

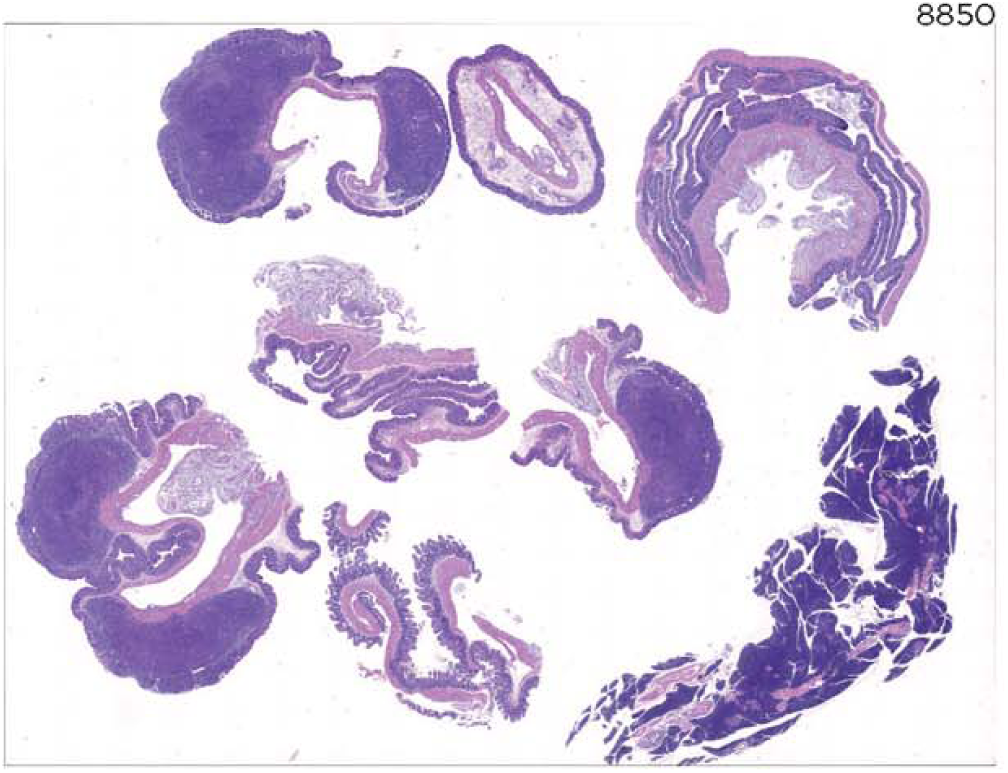

