## Supplemental Figures for "Disrupting ß-catenin dependent Wnt signaling activates an invasive gene program predictive of colon cancer progression"

**Supplementary Materials for Chen et al:**

Key for orthotopic tumor sections used in pathology scoring

| Mouse ID | Cell Line | Expression Vector |
|---|---|---|
| 13-38 | SW620 | Mock |
| 13-39 | SW620 | dnLEF1 |
| 14-01 | SW480 | Mock |
| 14-02 | SW480 | dnLEF1 |
| 8760 | SW620 | dnLEF1 |
| 8761 | SW620 | Mock |
| 8765 | SW620 | Mock |
| 8767 | SW620 | dnLEF1 |
| 8768 | SW620 | dnLEF1 |
| 8721 | COLO320 | LRP6KO |
| 8772 | SW620 | LRP6KO |
| 8777 | SW620 | Cas9 |
| 8778 | SW620 | Cas9 |
| 8803 | SW620 | LRP6KO |
| 8804 | SW620 | LRP6KO |
| 8805 | SW620 | Cas9 |
| 8806 | SW620 | Cas9 |
| 8791 | SW620 | Cas9 |
| 8793 | SW620 | LRP6KO |
| 8823 | COLO320 | LRP6KO |
| 8840 | COLO320 | Cas9 |
| 8788 | SW620 | LRP6KO |
| 8790 | SW620 | Cas9 |
| 8773 | SW620 | Mock |
| 8774 | SW620 | dnLEF1 |
| 8775 | SW620 | dnLEF1 |
| 8782 | SW620 | Mock |
| 8830 | SW480 | Mock |
| 8831 | SW480 | Mock |
| 8835 | SW480 | dnLEF1 |
| 8836 | SW480 | dnLEF1 |
| 8837 | SW480 | dnLEF1 |
| 8839 | SW480 | Mock |
| 8825 | COLO320 | LRP6KO |
| 8827 | COLO320 | Cas9 |
| 8849 | COLO320 | Cas9 |
| 8850 | COLO320 | LRP6KO |

13-38

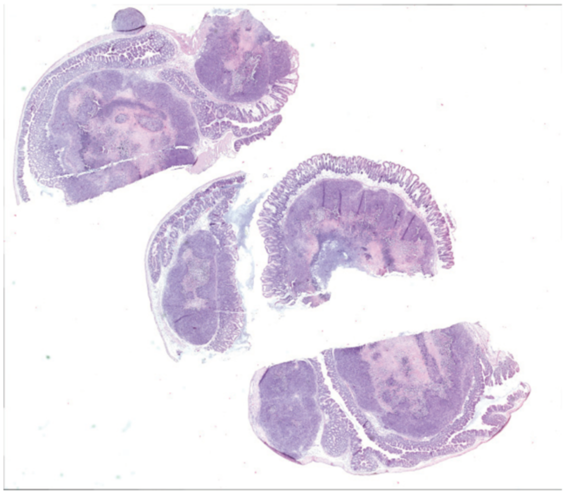

13-39

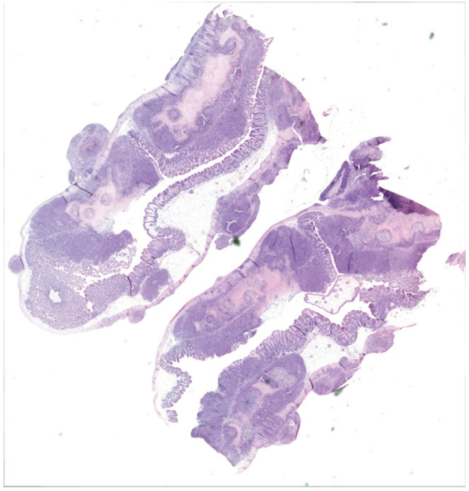

14-01

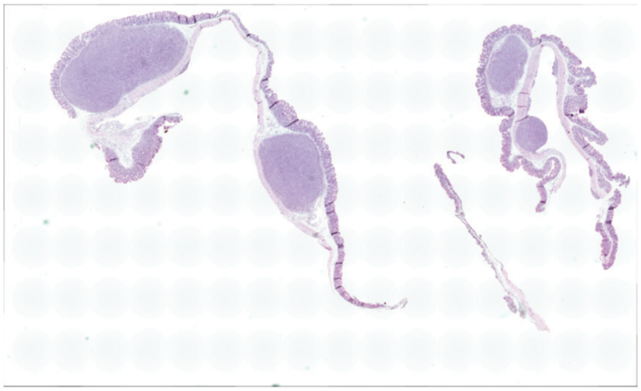

14-02

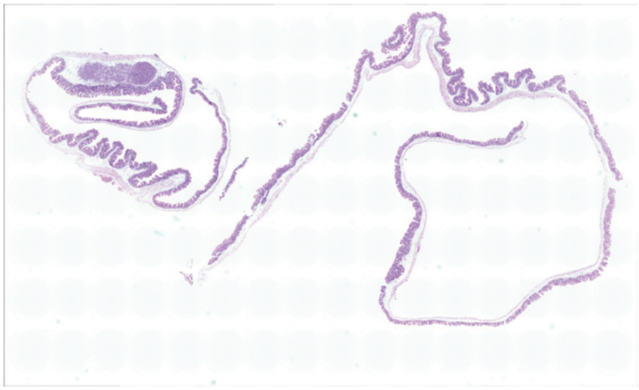

8760

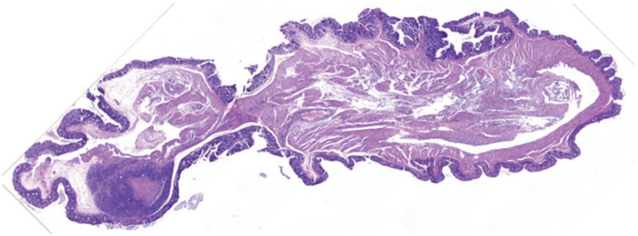

8761

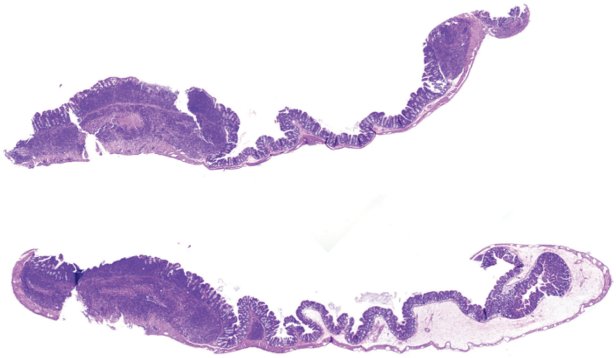

8765

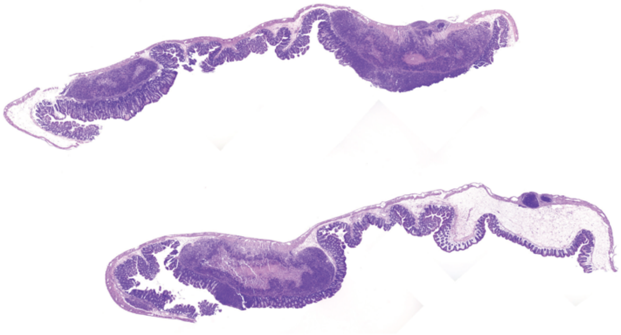

8767

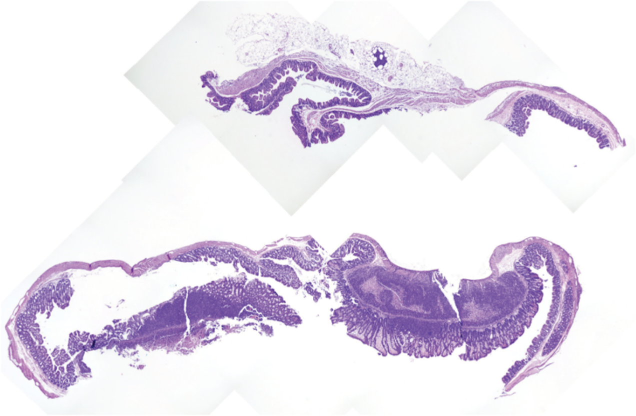

8768

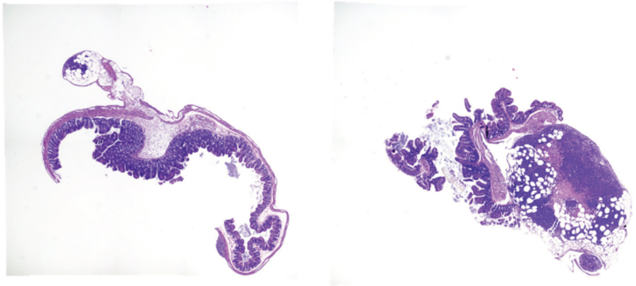

8721

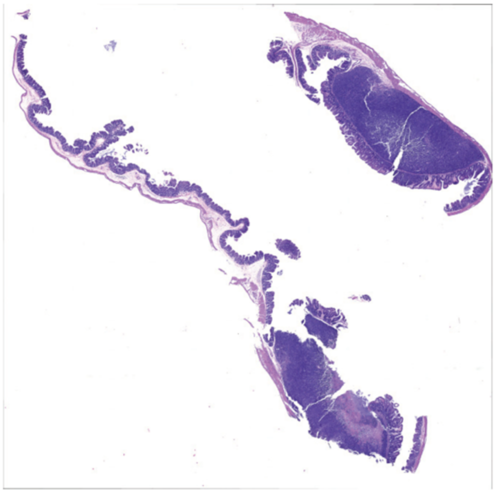

8772

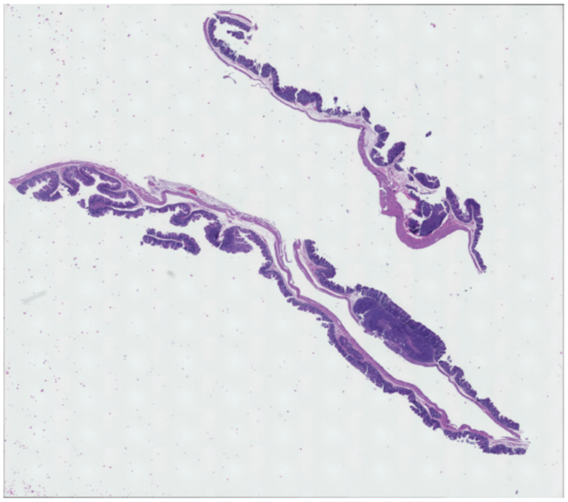

8777

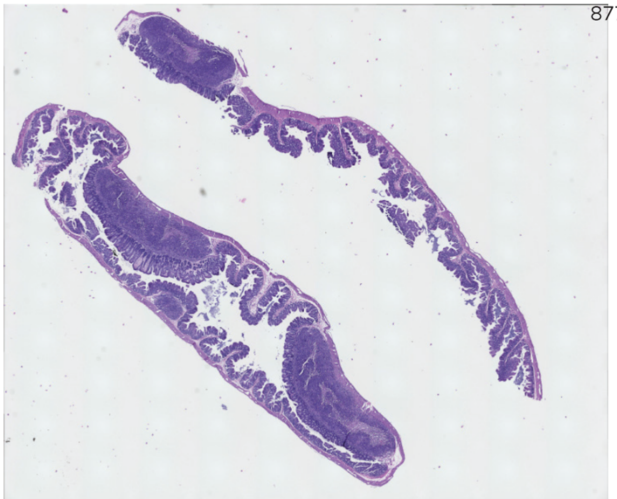

8778

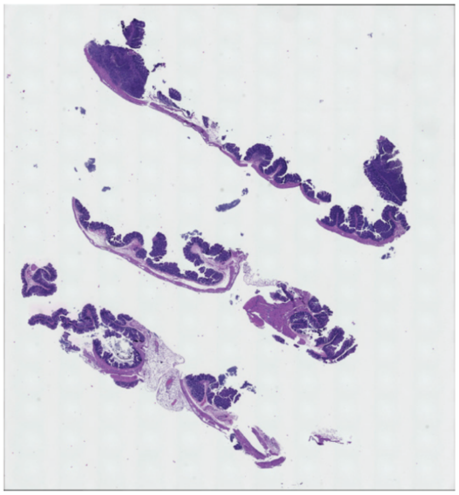

8803

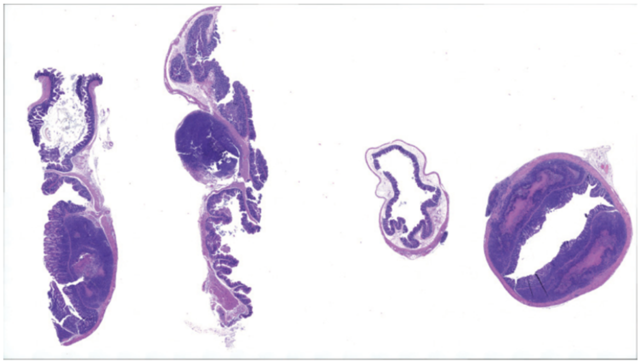

8804

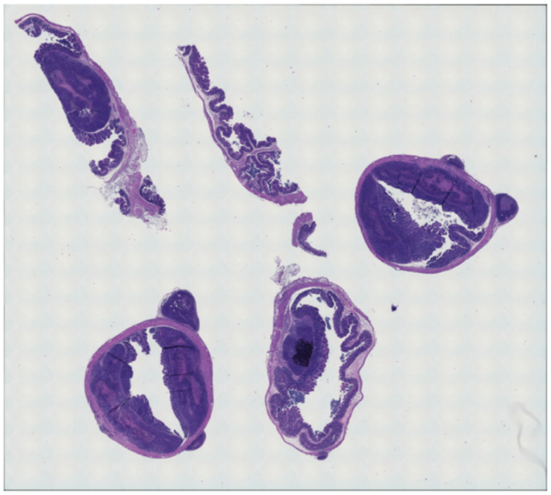

8805

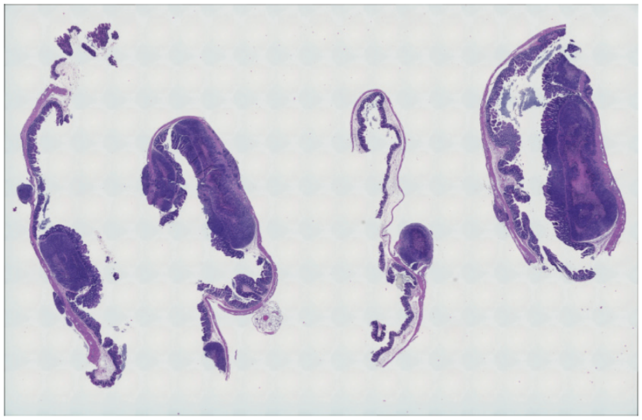

8806

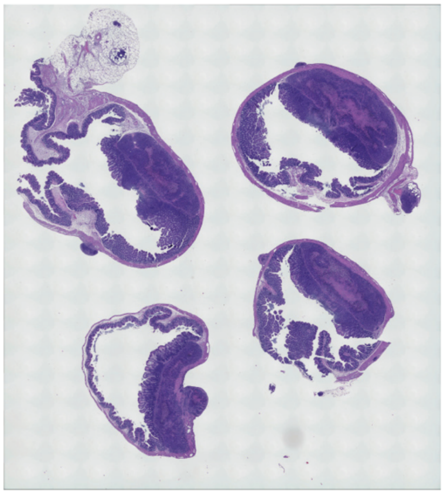

8791

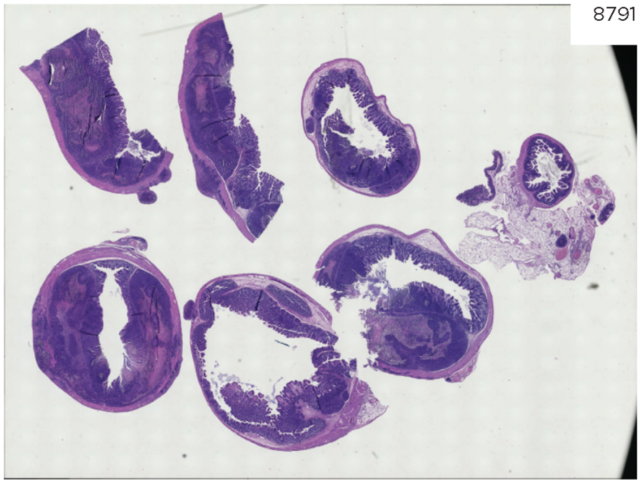

8793

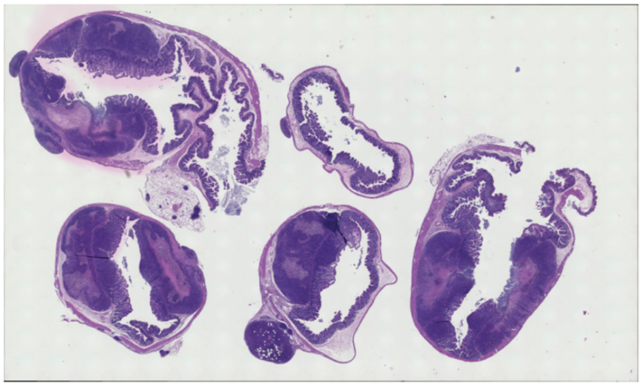

8823

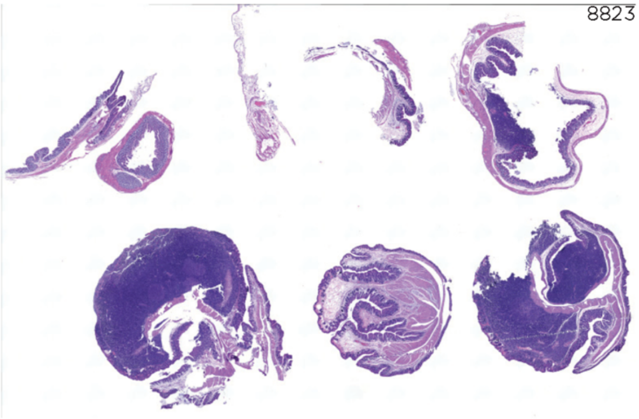

8840

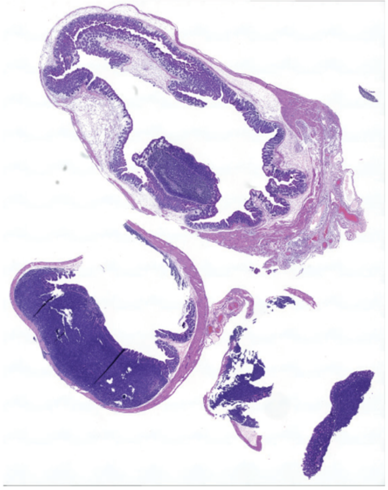

8788

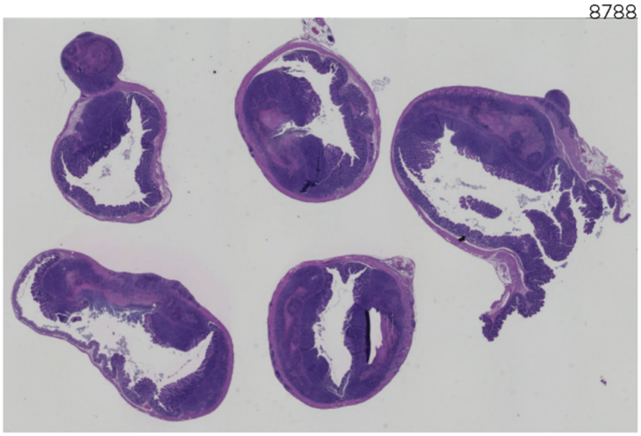

8790

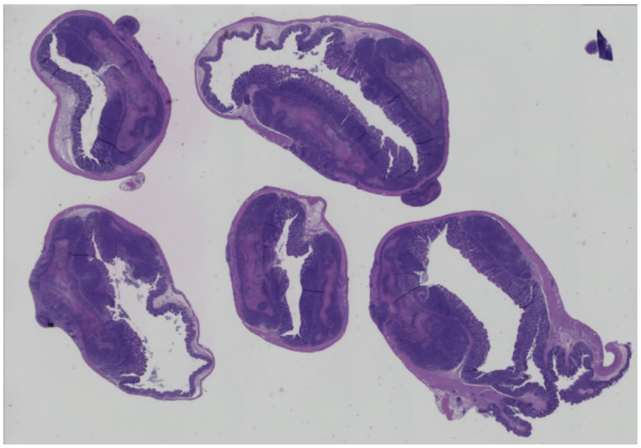

8773

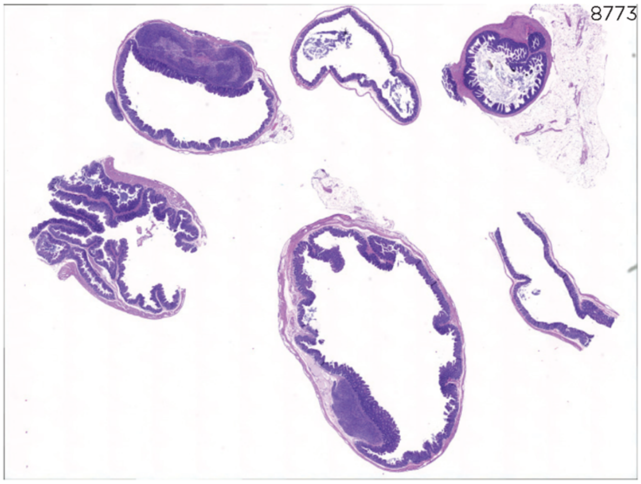

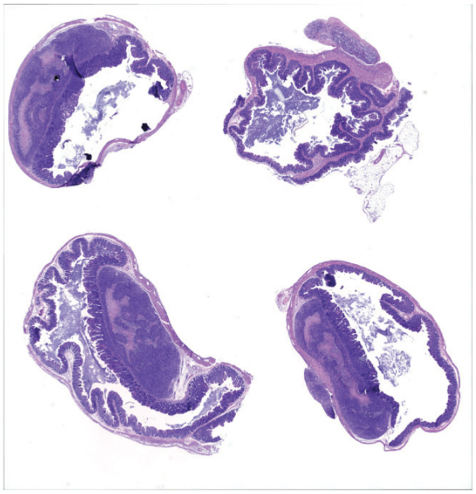
8774

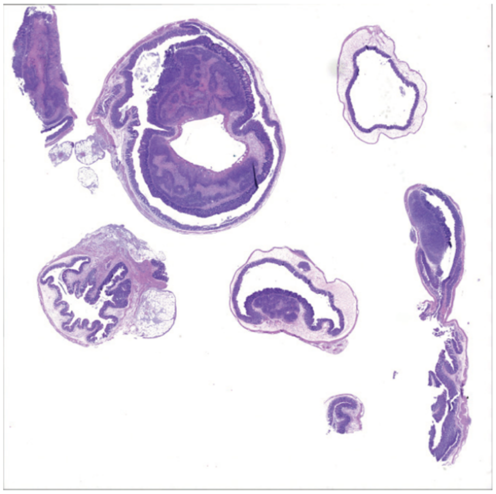
8775

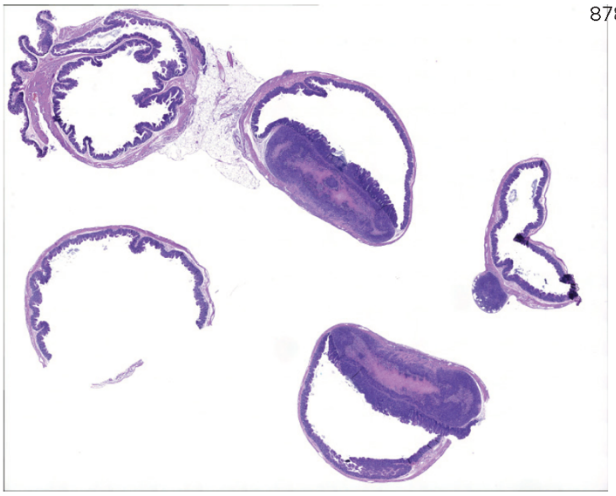
8782

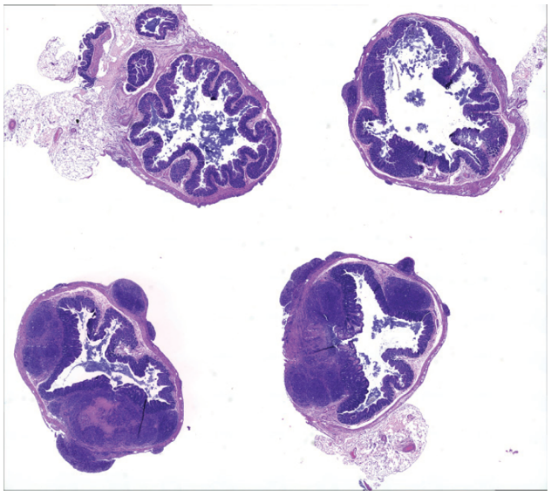
8830

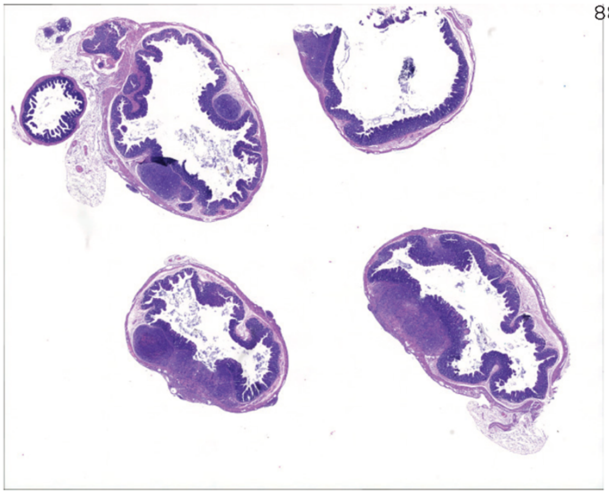
8831

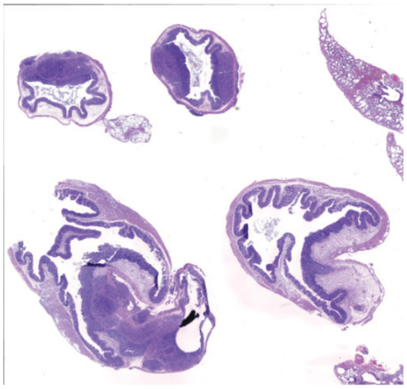
8835

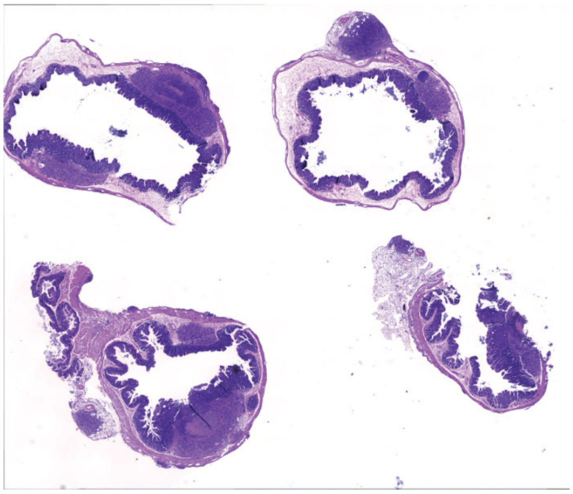
8836

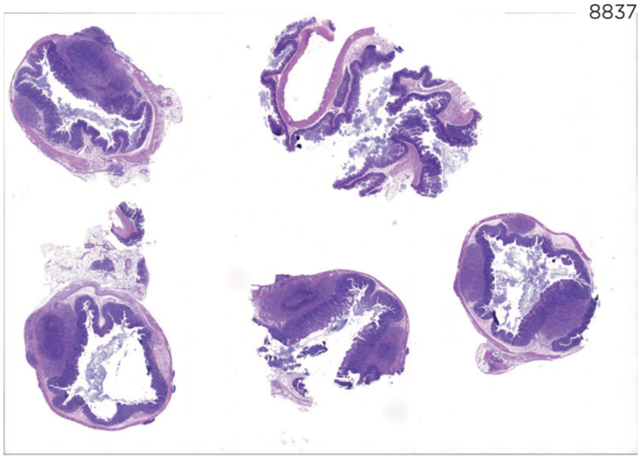
8837

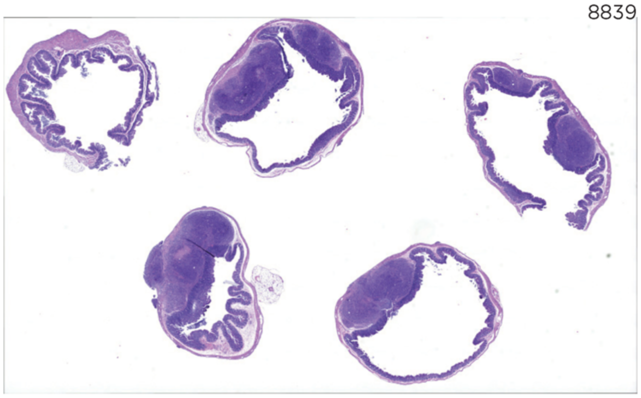
8839

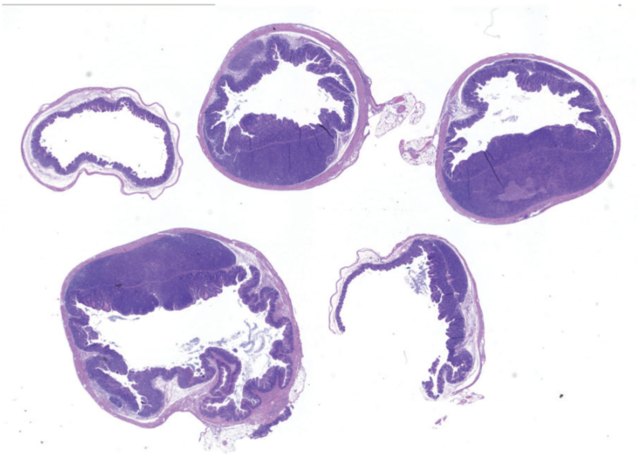
8825

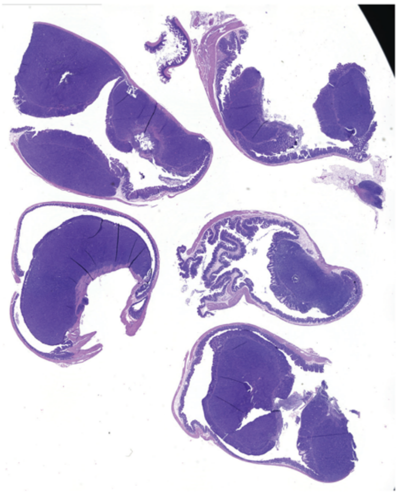
8827

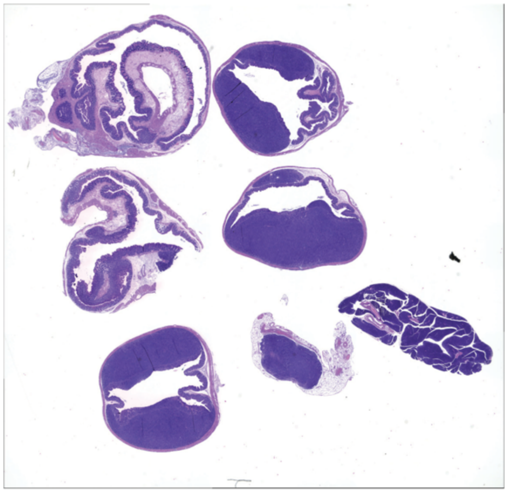
8849

8850

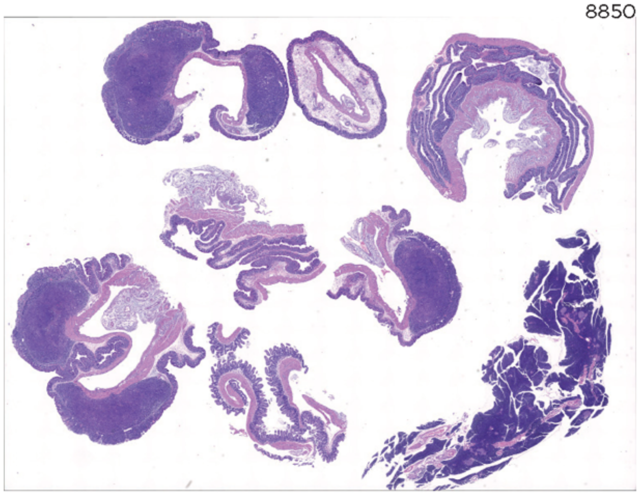

**Supplemental Figure 1. Orthotopic tumor sections for blinded pathology scoring.**

A. SW480 Mock & SW480 dnLEF1

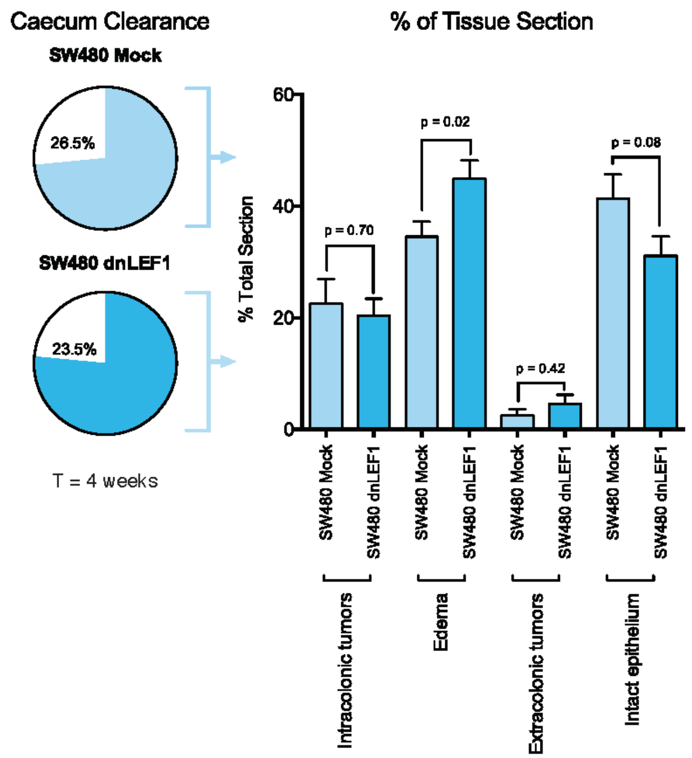


B. SW620 Mock & SW480 dnLEF1

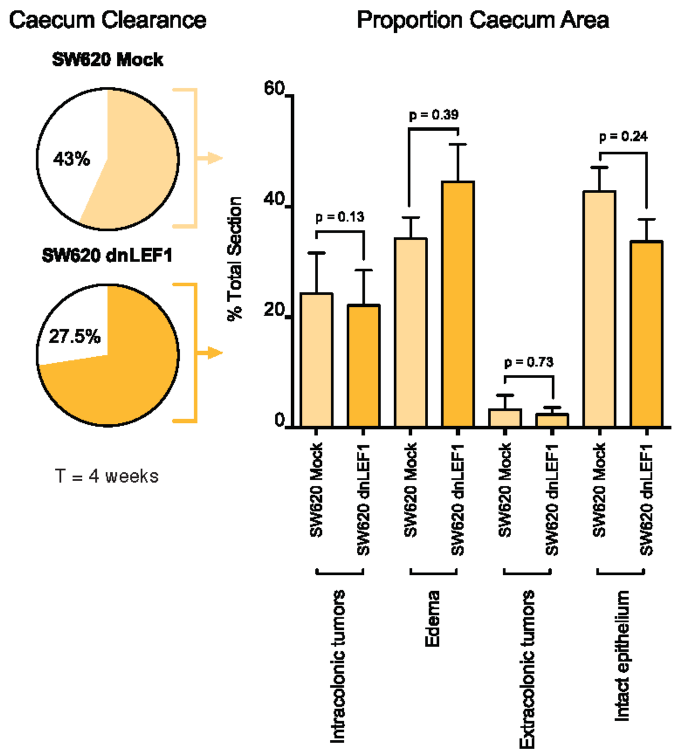


C. SW620 Cas9 & SW620 LRP6KO

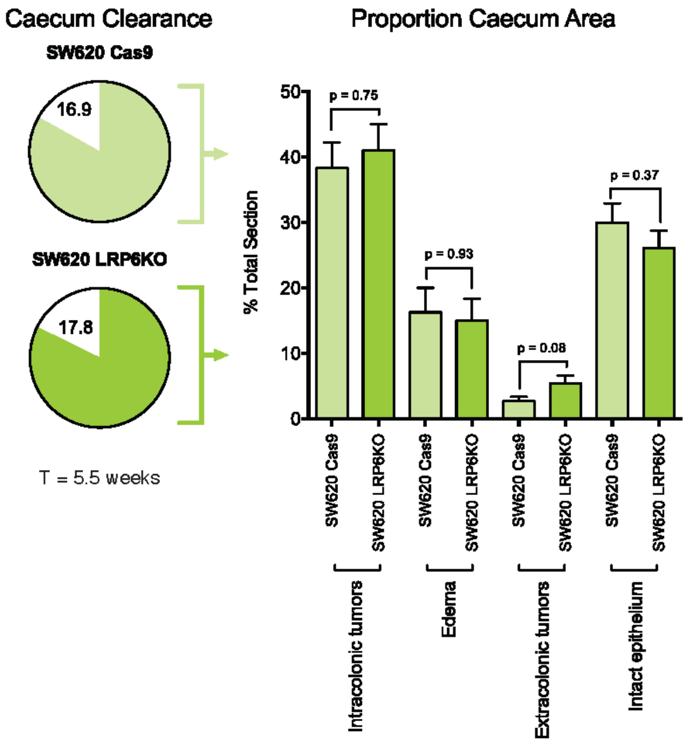

**Supplemental Figure 2. Quantification of orthotopic tumors.**

A. Percent of lumen clearance shows no significant difference between SW480 Mock and SW480 dnLEF1. A significant increase in edema and a decrease in intact epithelia are seen in SW480 dnLEF1. B. Percent of lumen clearance shows a significant decrease in cleared area in SW620 dnLEF1 compared to SW620 Mock. An increase in edema and a decrease of intact epithelia are seen in SW480 dnLEF1, but not to significance. C. No significant difference in lumen clearance observed in SW620 LRP6KO, but an increase in extracolonic tumors is observed.

A. LRP5 CRISPR

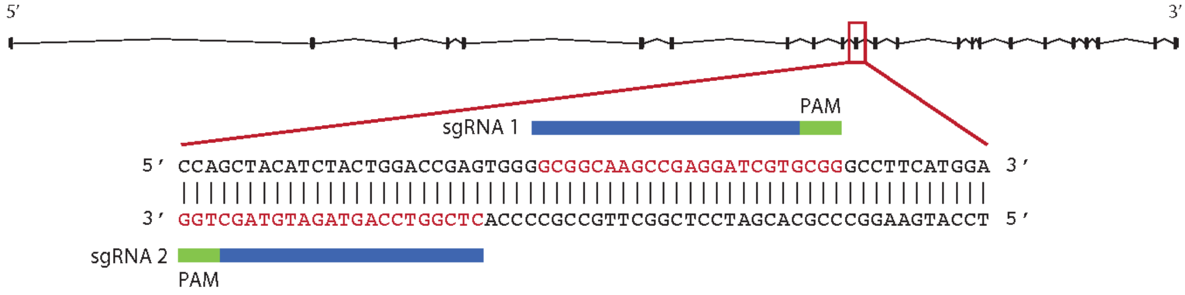


B.

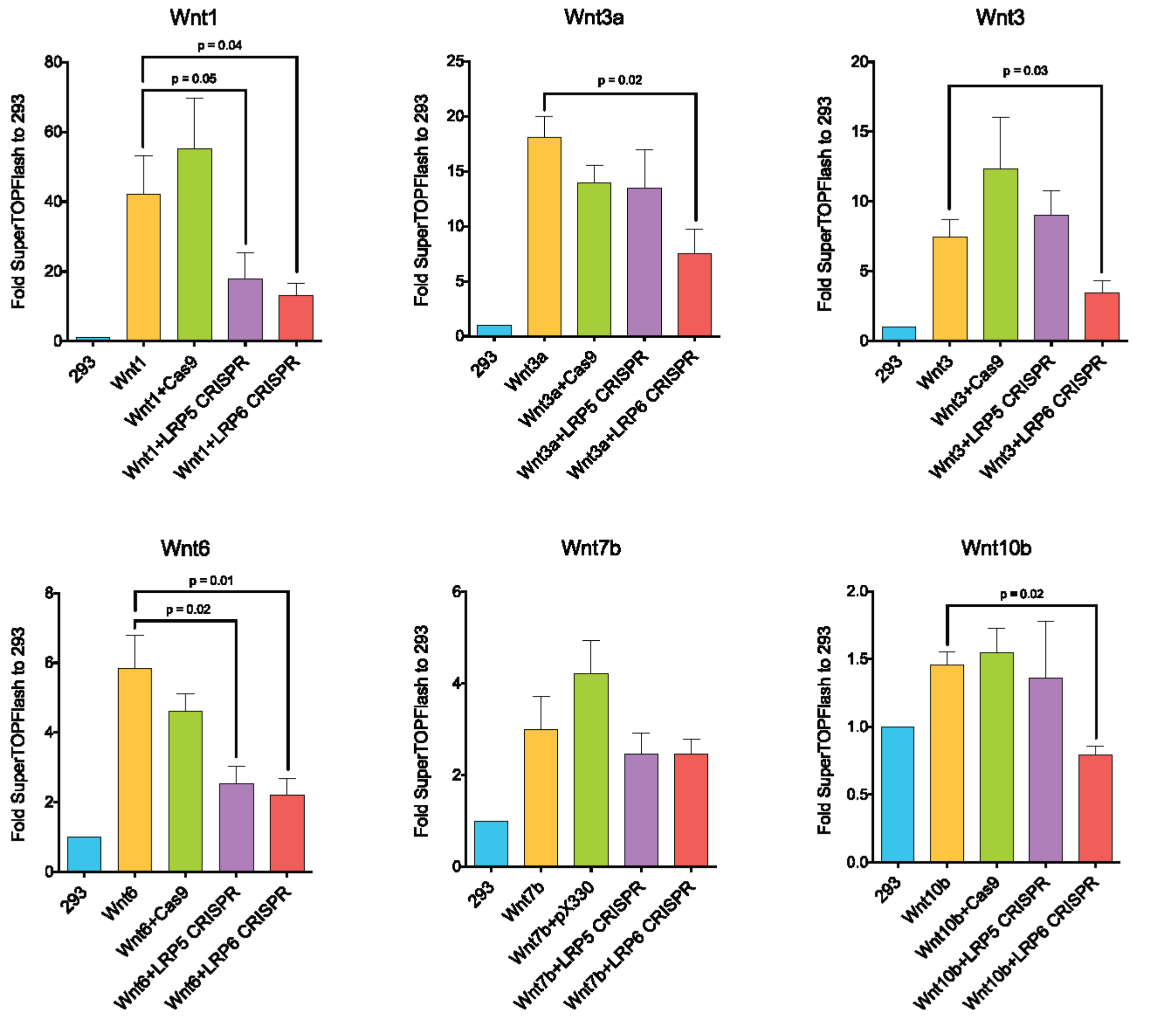


N = 4, SEM shown

C.

| | Wnt1 | Wnt3a | Wnt3 | Wnt6 | Wnt7b | Wnt10b |
|---|---|---|---|---|---|---|
| LRP5 | ✓ | | | ✓ | | |
| LRP6 | ✓ | ✓ | ✓ | ✓ | | ✓ |

Required for signaling

**Supplemental Figure 3. Wnt ligands require LRP6 to activate Wnt signaling.**

A. LRP5 CRISPR construct. B. HEK293 cells were transiently transfected with a Wnt ligand and a LRP5/6 CRISPR construct. Of the ligands tested, all except Wnt7b showed a significant decrease in Wnt signaling activity with the loss of LRP6. A number of Wnt ligands also showed sensitivity to loss of LRP5, however, no Wnt ligand was solely active through LRP5. Additionally, loss of LRP5 or LRP6 decreased Wnt activity by more than 50%, suggesting that HEK293 cells were not simply switching reliance to the other LRP co-receptor. We find that these ligands require both LRP5 and LRP6 to activate signaling. C. Summary of Wnt ligand requirements for LRP5 and LRP6.

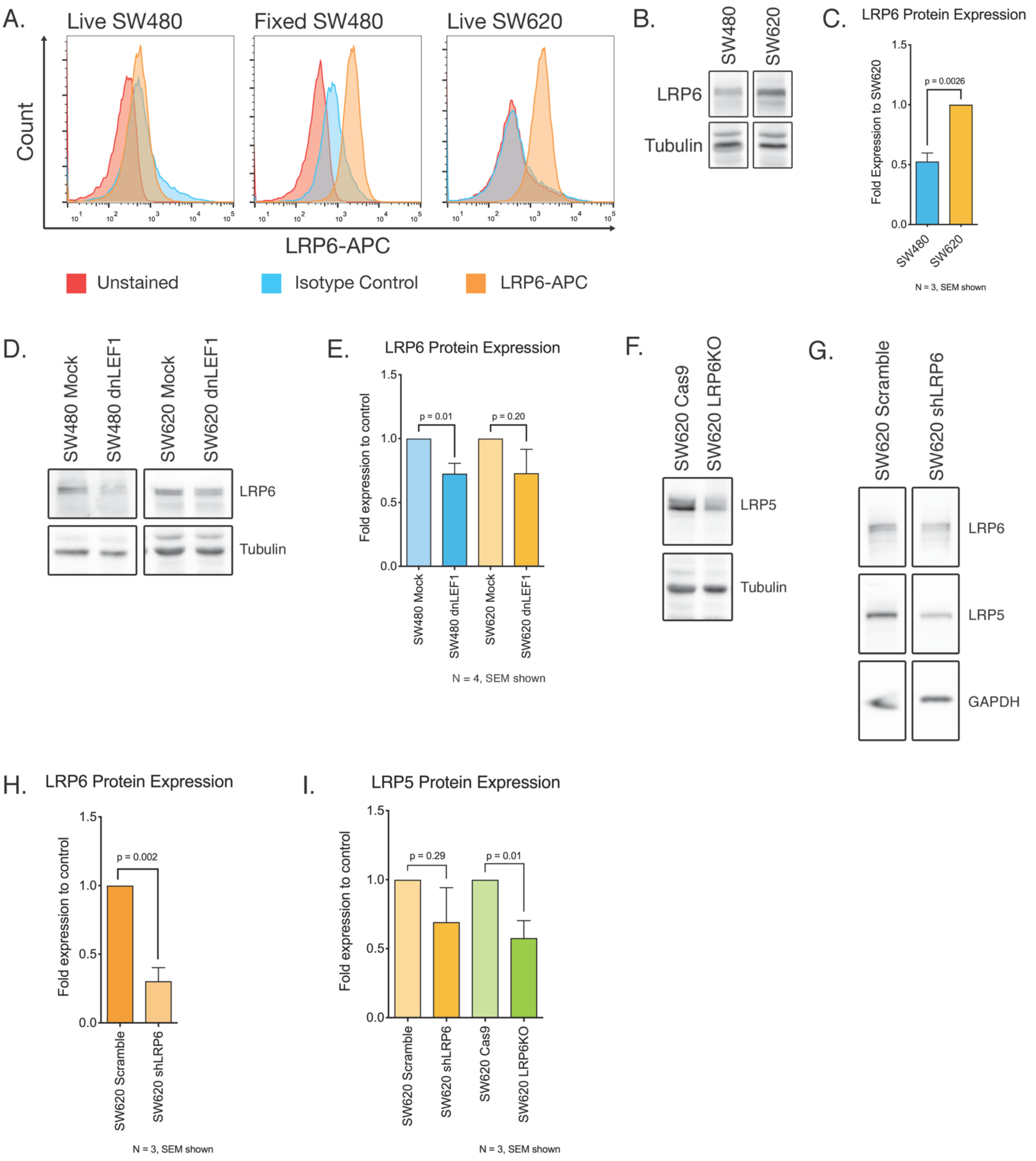
A.
Live SW480
Fixed SW480
Live SW620
Count
LRP6-APC
Unstained
Isotype Control
LRP6-APC
B.
SW480
SW620
LRP6
Tubulin
C.
LRP6 Protein Expression
Fold Expression to SW620
p = 0.0026
SW480
SW620
N = 3, SEM shown
D.
SW480 Mock
SW480 dnLEF1
SW620 Mock
SW620 dnLEF1
LRP6
Tubulin
E.
LRP6 Protein Expression
Fold expression to control
p = 0.01
p = 0.20
SW480 Mock
SW480 dnLEF1
SW620 Mock
SW620 dnLEF1
N = 4, SEM shown
F.
SW620 Cas9
SW620 LRP6KO
LRP5
Tubulin
G.
SW620 Scramble
SW620 shLRP6
LRP6
LRP5
GAPDH
H.
LRP6 Protein Expression
Fold expression to control
p = 0.002
SW620 Scramble
SW620 shLRP6
N = 3, SEM shown
I.
LRP5 Protein Expression
Fold expression to control
p = 0.29
p = 0.01
SW620 Scramble
SW620 shLRP6
SW620 Cas9
SW620 LRP6KO
N = 3, SEM shown

**Supplemental Figure 4. LRP6 is not expressed on the cell membrane of SW480 cells**

A. Flow cytometry of SW480 and SW620 cells reveals that LRP6 is not detectable in SW480 cells, but a signal is clearly detected after cells were fixed and permeabilized, indicating LRP6 protein is present but not localized to the cell membrane. B-C. Western blot of whole cell lysates from SW480 and SW620 cells probed for LRP6 show that SW620 cells express significantly more LRP6 protein than SW480 cells. D-E. LRP6 protein expression is decreased in dnLEF1-expressing cell lines

Total LRP6 protein is significantly decreased in SW480 dnLEF1 cells compared to Mock. A noticeable but not significant decrease in LRP6 is observed in SW620 dnLEF1 cells. F-H. Loss of LRP6 induces a loss of LRP5 in SW620 cells

F. By Western blotting, we find a significant decrease in LRP5 protein in LRP6 knockout cells. To confirm that this was not an off-target effect of our construct, we used a lentiviral shRNA to knockout LRP6. Total LRP5 was also decreased in the shRNA-transduced cells.

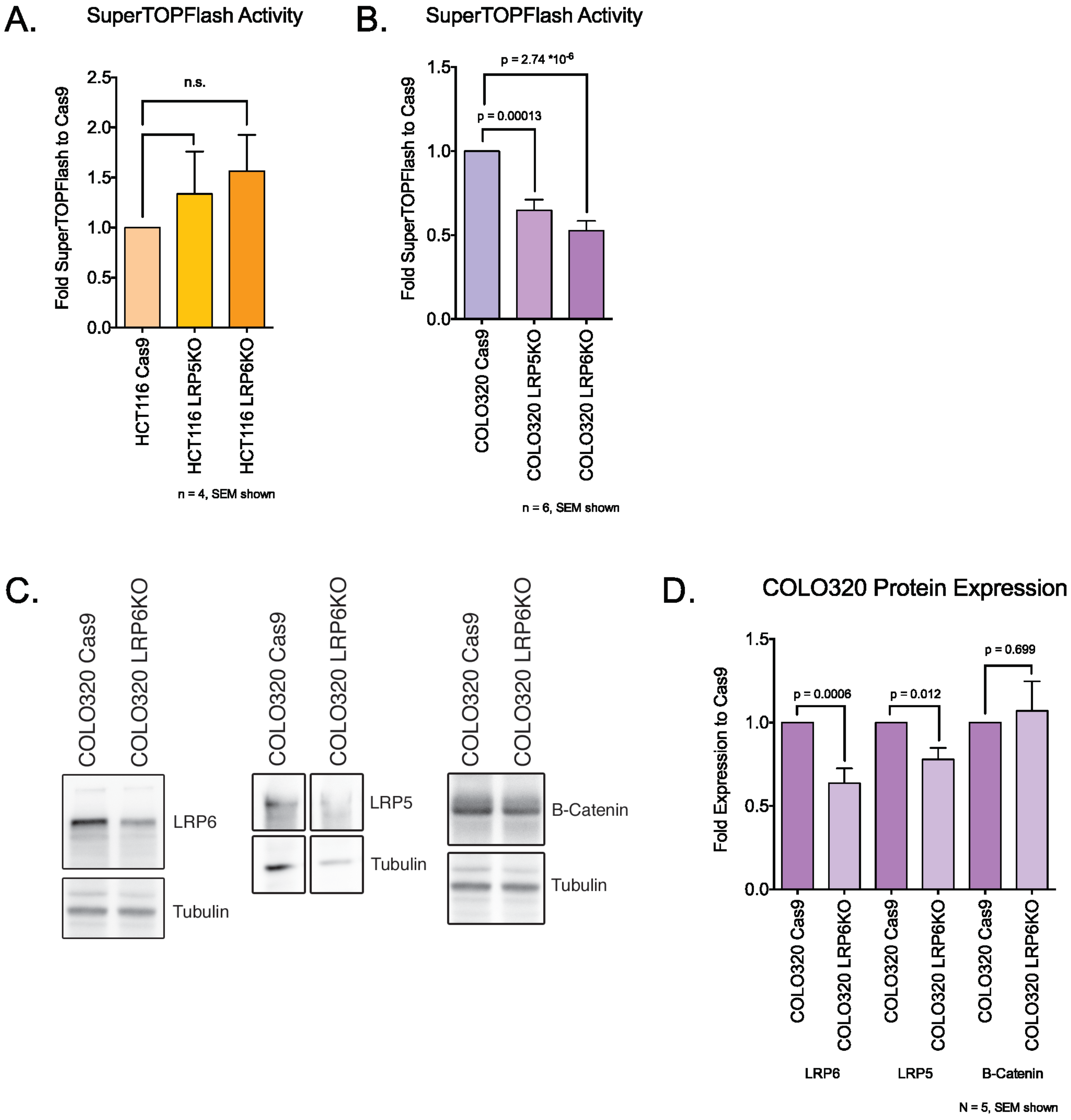
A.
SuperTOPFlash Activity
Fold SuperTOPFlash to Cas9
n.s.
HCT116 Cas9
HCT116 LRP5KO
HCT116 LRP6KO
n = 4, SEM shown
B.
SuperTOPFlash Activity
Fold SuperTOPFlash to Cas9
p = 2.74 *10-6
p = 0.00013
COLO320 Cas9
COLO320 LRP5KO
COLO320 LRP6KO
n = 6, SEM shown
C.
COLO320 Cas9
COLO320 LRP6KO
LRP6
Tubulin
LRP5
Tubulin
B-Catenin
Tubulin
D.
COLO320 Protein Expression
Fold Expression to Cas9
p = 0.0006
p = 0.012
p = 0.699
COLO320 Cas9
COLO320 LRP6KO
LRP6
LRP5
B-Catenin
N = 5, SEM shown

**Supplemental Figure 5. Loss of LRP6 reduces Wnt signaling in COLO320, but not HCT116 cells**

A. HCT116 cells do not show a significant change to Wnt signaling activity in response to loss of LRP5 or LRP6. HCT116 cells have a mutation in b-catenin that allows the cells to evade the destruction complex in order to activate Wnt target genes. This suggests that in HCT116 cells, Wnt ligands may play a negligible role in modulating signaling activity. B. COLO320 cells show a significant decrease in Wnt signaling activity with the loss of LRP5 or LRP6. C-D. Western blots and quantitation of COLO320 Cas9 and LRP6KO cells shows a decrease in total LRP6 protein as expected. Interestingly, a loss of LRP5 is also observed with the knockout of LRP6. This is also observed in SW620 cells (See Supplemental Figure 4). Unlike the SW620 cells, we do not find a significant decrease in b-catenin in the COLO320 LRP6KO cells, suggesting a different mode of Wnt signaling regulation.

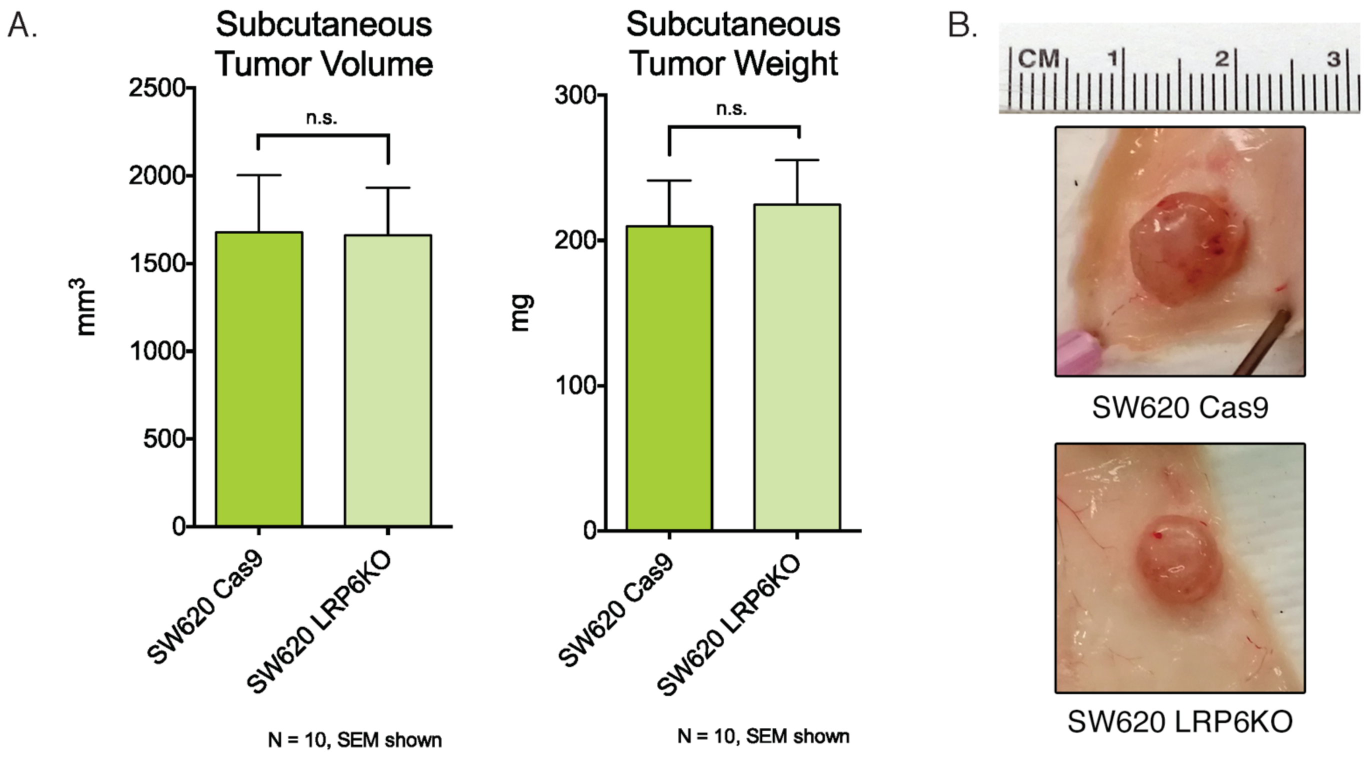
A.
Subcutaneous Tumor Volume
2500
2000
1500
1000
500
0
mm3
n.s.
SW620 Cas9
SW620 LRP6KO
N = 10, SEM shown
Subcutaneous Tumor Weight
300
200
100
0
mg
n.s.
SW620 Cas9
SW620 LRP6KO
N = 10, SEM shown
B.
CM 1 2 3
SW620 Cas9
SW620 LRP6KO

**Supplemental Figure 6. Subcutaneous tumors of SW620 Cas9 and LRP6KO**

A. No significant difference in tumor volume and weight is seen between SW620 Cas9 and SW620 LRP6KO. B. Representative images of tumors at harvest.

| Study | Group | Patients | % Gender | Age |
|---|---|---|---|---|
| GSE15781 | No Treatment | 11 | 54 M/46 F | 74 |
| | Treated | 10 | 80 M/20 F | 68.3 |
| GSE17536 | Total | 177 | 54 M/46 F | 65.5 ± 13.1 |
| GSE39582 | Total | 586 | 54.9 M/45.1 F | 66.95 ± 13.17 |

**Supplemental Figure 7. Patient characteristics from studies used in Kaplan-Meier curves** (figure 4).

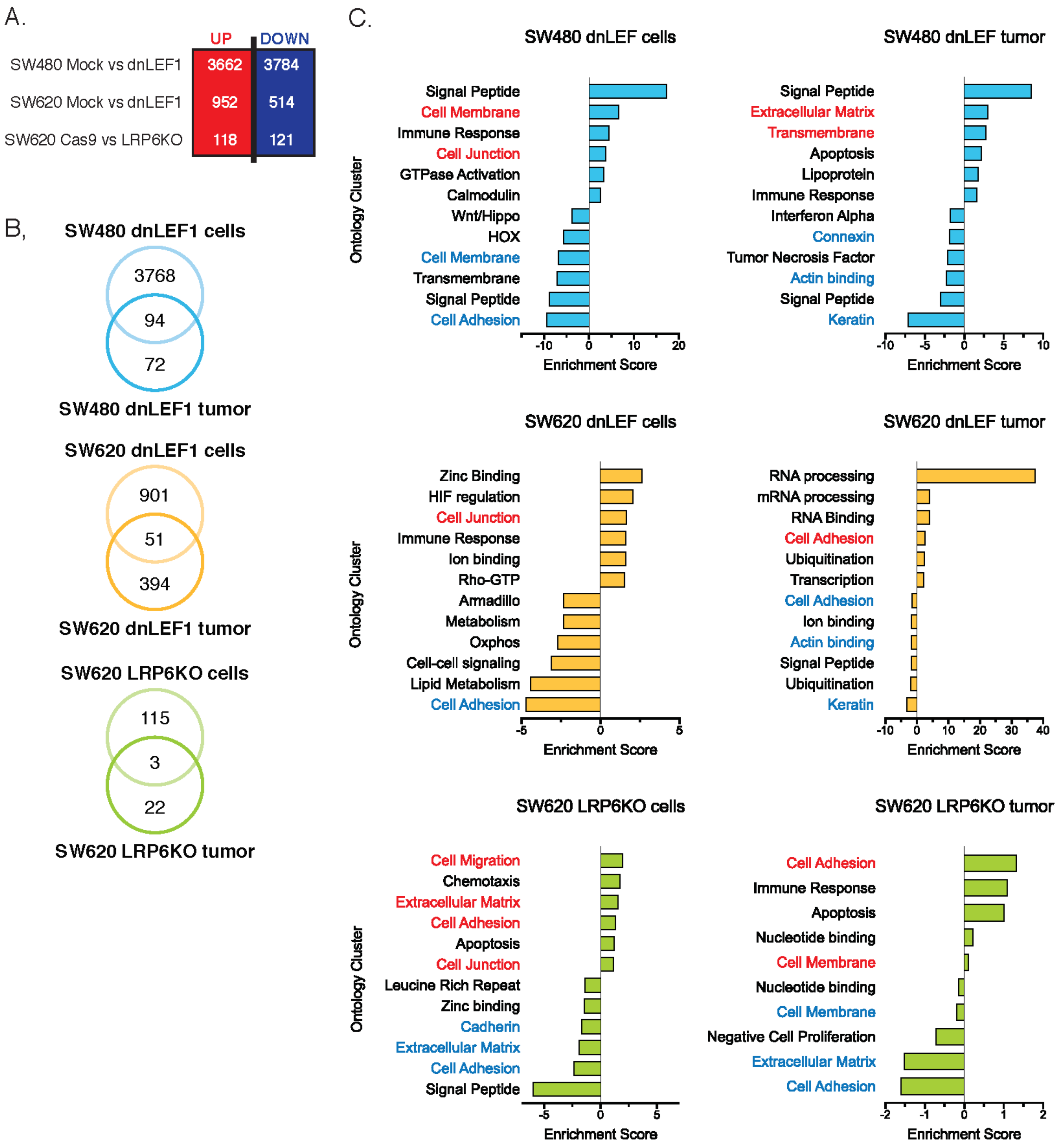
A.
UP
DOWN
SW480 Mock vs dnLEF1
3662
3784
SW620 Mock vs dnLEF1
952
514
SW620 Cas9 vs LRP6KO
118
121
B,
SW480 dnLEF1 cells
3768
94
72
SW480 dnLEF1 tumor
SW620 dnLEF1 cells
901
51
394
SW620 dnLEF1 tumor
SW620 LRP6KO cells
115
3
22
SW620 LRP6KO tumor
C.
SW480 dnLEF cells
Ontology Cluster
Signal Peptide
Cell Membrane
Immune Response
Cell Junction
GTPase Activation
Calmodulin
Wnt/Hippo
HOX
Cell Membrane
Transmembrane
Signal Peptide
Cell Adhesion
-10
0
10
20
Enrichment Score
SW480 dnLEF tumor
Signal Peptide
Extracellular Matrix
Transmembrane
Apoptosis
Lipoprotein
Immune Response
Interferon Alpha
Connexin
Tumor Necrosis Factor
Actin binding
Signal Peptide
Keratin
-10
-5
0
5
10
Enrichment Score
SW620 dnLEF cells
Ontology Cluster
Zinc Binding
HIF regulation
Cell Junction
Immune Response
Ion binding
Rho-GTP
Armadillo
Metabolism
Oxphos
Cell-cell signaling
Lipid Metabolism
Cell Adhesion
-5
0
5
Enrichment Score
SW620 dnLEF tumor
RNA processing
mRNA processing
RNA Binding
Cell Adhesion
Ubiquitination
Transcription
Cell Adhesion
Ion binding
Actin binding
Signal Peptide
Ubiquitination
Keratin
-10
0
10
20
30
40
Enrichment Score
SW620 LRP6KO cells
Ontology Cluster
Cell Migration
Chemotaxis
Extracellular Matrix
Cell Adhesion
Apoptosis
Cell Junction
Leucine Rich Repeat
Zinc binding
Cadherin
Extracellular Matrix
Cell Adhesion
Signal Peptide
-5
0
5
Enrichment Score
SW620 LRP6KO tumor
Cell Adhesion
Immune Response
Apoptosis
Nucleotide binding
Cell Membrane
Nucleotide binding
Cell Membrane
Negative Cell Proliferation
Extracellular Matrix
Cell Adhesion
-2
-1
0
1
2
Enrichment Score

**Supplemental Figure 8. Decreased Wnt signaling induces global transcriptome changes *in vitro* in genes associated with cell adhesion and extracellular matrix.** A. Significantly changed genes between Wnt-decreased cell lines and their associated control. B. Overlap of significantly upregulated genes between RNA from cell culture and RNA from orthotopic tumors shows relatively few conserved gene expression changes. C. Top DAVID gene ontology clusters for significantly upregulated and downregulated genes *in vitro* and *in vivo* reveal conserved changes in gene programs associated with cell adhesion and extracellular matrix.

Data files S1-S#
